# PSMε controls skin commensal CD8^+^ T cell activation

**DOI:** 10.64898/2026.07.30.741773

**Authors:** William S. Owens, Kevin Lenzi, Wendy Geng, Annalisa Hurd, Jessie J. Y. Liu, Caleb Staudinger, Brittany Berdy, Cameron Lian, Namrata D. Udeshi, Steven A. Carr, Christopher D. Johnston, Jonathan Livny, Y. Erin Chen

## Abstract

Upon skin colonization, the prevalent human skin commensal *S. epidermidis* can elicit a CD8^+^ T cell response that protects against pathogens or clears tumors. The microbial features that drive this response are undefined, limiting our ability to understand and predict commensal-immune crosstalk and to engineer potent commensal-derived immunotherapies. To uncover these microbial features, we harnessed both the natural variation in CD8^+^ T cell induction across primary human isolates of *Staphylococcus* and our ability to genetically manipulate these strains. Stimulatory strains exhibit increased quorum sensing activation, which turns on a unique commensal-associated gene family, called phenol-soluble modulin ε (PSMε), that is required for CD8^+^ T cell activation. PSMε not only acts as the immunodominant CD8^+^ T cell antigen but also enhances cross-presentation in an antigen-agnostic manner. Co-delivering PSMε promotes CD8^+^ T cell priming to an exogenous antigen via a mechanism that is independent of formyl peptide receptor and co-stimulatory receptor upregulation. Thus, we demonstrate that specific aspects of microbiome-immune crosstalk can be distilled to molecular components, which engage in previously undescribed mechanisms and can be harnessed for immunotherapy without requiring live bacterial colonization.

## MAIN TEXT

Some members of our commensal microbiota prime and activate potent antigen-specific T cells upon skin or gut colonization^1–6^. These T cells modulate diverse processes, including pathogen resistance, wound healing, and antitumor immunity^1,7–10^. We have a growing appreciation that the host mechanisms underlying these commensal T cell responses are distinct from those governing infection-induced T cell responses. For example, epithelial cells, rather than professional immune cells, can present antigens directly to gut commensal-specific CD8^+^ T cells^11^ and epithelial upregulation of endogenous retroviruses promotes the expansion of skin commensal-specific CD8^+^ T cells^12^. However, the *microbial* mechanisms underlying commensal-specific T cell responses remain largely uncharacterized.

What distinguishes a bacterial strain that potently activates certain T cell subsets from closely related non-activating strains is often unknown. Because related commensals can colonize with similar abundance, do not cause infection at homeostasis, and do not differ in canonical virulence factors, the key molecular features that distinguish immune-stimulatory from non-stimulatory strains are not obvious. This fundamental gap in knowledge prevents the field from converting microbiome sequencing data into functional predictions of host immune responses and from effectively harnessing specific components of our microbiome as therapeutics.

We sought to address this gap for an unusual and useful commensal T cell response: the CD8^+^ T cell response elicited by the prevalent human skin commensal *Staphylococcus epidermidis* (*S. epi*)^1,7^. We previously showed that skin colonization by tumor antigen-expressing *S. epi* activates tumor-specific CD8^+^ T cells, which protect mice from both skin and metastatic melanoma^13^. This finding that cytotoxic CD8^+^ T cells could be elicited across an intact epithelial barrier without innate immune inflammation was surprising and opened the door to developing highly targeted, needle-free immunotherapies using commensal bacterial components. Yet, the underlying microbial mechanisms were unknown. While many commensals can induce CD4^+^ T cells^2–4,6,14–16^, relatively few commensals are known to induce CD8^+^ T cells like *S. epi*^10^. Even among *S. epi* strains, only a subset can activate CD8^+^ T cells, despite their genomic similarity, suggesting that simply colonizing the skin and expressing canonical innate immune ligands, such as peptidoglycan or wall teichoic acids, is not sufficient^8^. Although we previously found that certain pattern recognition receptors (PRRs), including TLR2, CLEC7A, and cGAS-STING, are necessary for the *S. epi* CD8^+^ T cell response^12,17^, they are not sufficient to distinguish between CD8^+^ T cell-activating (i.e. “stimulatory”) versus non-stimulatory *Staphylococcus* strains.

Here, we define the genetic and molecular mechanisms that distinguish stimulatory strains of *Staphylococcus* from non-stimulatory strains; in doing so, we discovered an unexpected and novel microbial mechanism for promoting antigen cross-presentation. First, we found that stimulatory strains exhibit enhanced activation of a conserved microbe-microbe communication pathway called quorum sensing. Quorum sensing regulates a broad transcriptional program, but we found that one of its targets is critical for CD8^+^ T cell activation: phenol-soluble modulin epsilon (PSMε), a gene family unique to commensal *Staphylococcus*. PSMε has two key functions that can be decoupled. The first six amino acids make up the immunodominant epitope for *S. epi*-specific CD8^+^ T cells, while full-length PSMε enhances its own cross-presentation, a process required for most CD8^+^ T cell responses. Remarkably, purified PSMε is sufficient to enhance the cross-presentation of an exogenous co-delivered antigen. This function is unique to PSMε and not recapitulated by other *Staphylococcus* PSMs. Additionally, the requirement for quorum sensing activation and PSMε not only applies to *S. epi* but is conserved across *Staphylococcus* species; we demonstrate this requirement in detail for another human skin commensal *S. caprae*. Thus, PSMε is a unique commensal gene that is upregulated by quorum sensing activation and explains the ability of certain *Staphylococcus* strains to potently activate CD8^+^ T cells during physiologic skin colonization. PSMε does not act like a typical adjuvant that solely engages PRRs and innate immune signaling; rather, it specifically enhances antigen cross-presentation, suggesting high potential therapeutic utility.

### CD8^+^ T cell responses to skin commensals are restricted to specific *Staphylococcus* strains

Since *S. epi* NIHLM087 (LM087) elicits a potent antigen-specific CD8^+^ T cell response but many other strains of *S. epi* do not^8,13,18^, we first wanted to determine if CD8^+^ T cells are an unusual or common response to skin colonists. Thus, we screened across the three dominant genera of human skin commensal bacteria (*Staphylococcus*, *Corynebacterium*, and *Cutibacterium*) for their ability to elicit T cell responses upon skin colonization of conventionally housed C57BL/6 mice, without any barrier breach or additional skin preparation (**Figure 1A-F**). We particularly focused on screening diverse species of *Staphylococcus*, as this genus contains the only previously described CD8^+^ T cell activators among skin commensals^1,7,8^. Two weeks after the initial colonization, the skin (ears) and skin-draining cervical lymph nodes (cLN) were collected and analyzed for colonization burden (**Figure 1C**) and immune cell content (**Figure 1D-E**). Despite no major changes in total cLN cellularity, suggesting a lack of nonspecific inflammation (**Figure S1A**), almost all *Staphylococcus* strains expanded CD4^+^ effector T cells (Foxp3^−^ CD62L^-^ CD44^+^ Teff) and CCR6^+^ T cells (i.e. Th17-biased cells^19^ characteristic of extracellular bacterial responses) in both the cLN and ears (**Figure 1D**). These data suggest that most *Staphylococcus* strains are sensed by the adaptive immune system without barrier breach and that their antigens reach the lymph node. In contrast, only a few strains expanded CD8^+^ Teff and CCR6^+^ Tc17 cells in the cLN and skin, specifically two closely related *S. epi* isolates LM087 and NIHLM040 (LM040), a more distant *S. epi* isolate W23144, and *S. caprae* ATCC 55133 (*S. cap*) (**Figure 1E**). Importantly, these differences are not explained by colonization burden, which was comparable across all *Staphylococcus* and did not significantly correlate with skin expansion of CD8^+^ CCR6^+^ T cells (Tc17^8,18^) (**Figure 1C, F**). Overall, while CD4^+^ T cell responses to skin commensals are common, CD8^+^ T cell responses are rarer but not completely restricted to *S. epi* and can occur with other *Staphylococcus* species, such as *S. cap*.

**Figure 1.**
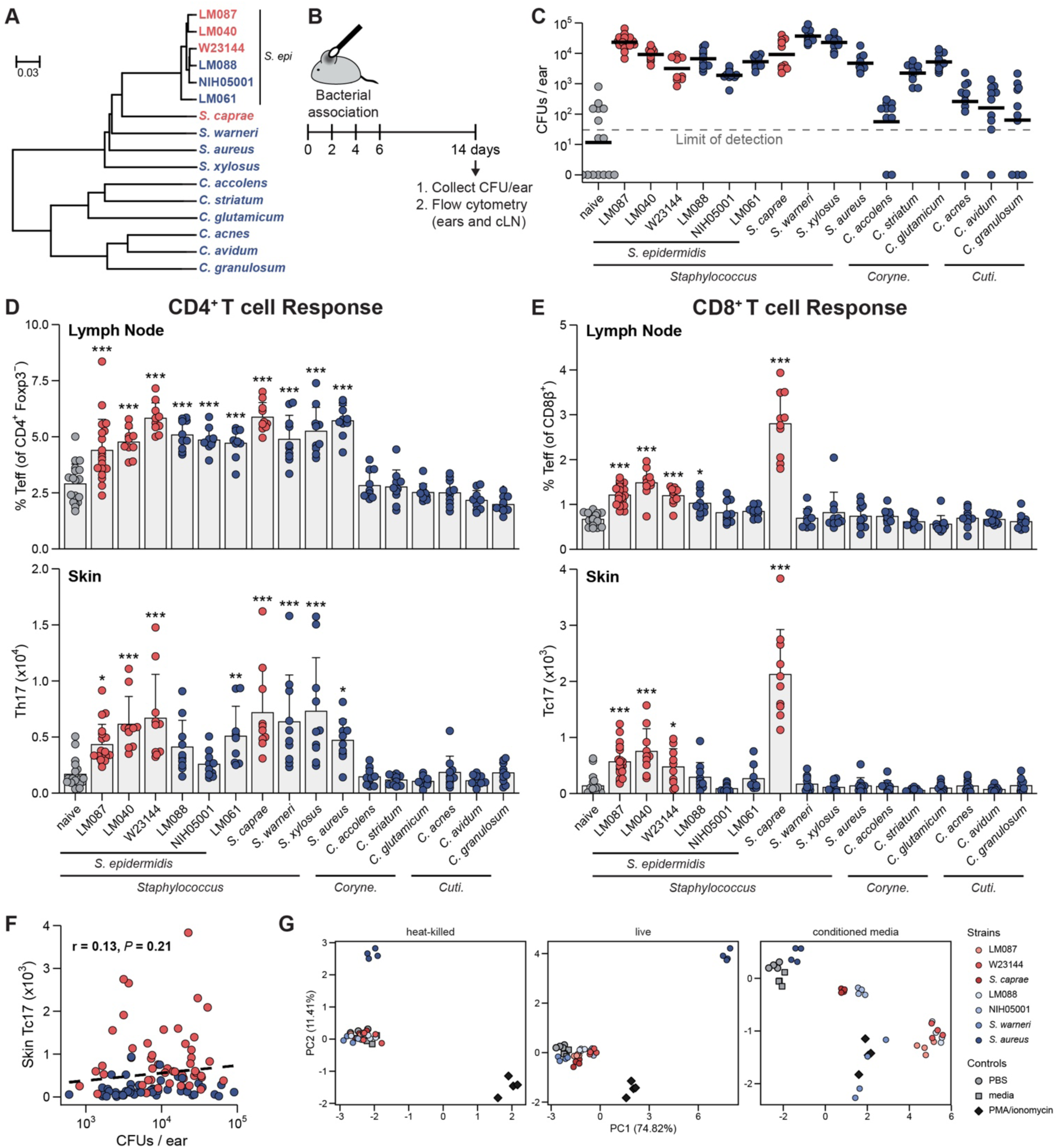
CD8^+^ T cell responses to skin commensals are restricted to specific *Staphylococcus* strains. (**A**) Whole-genome phylogenetic tree of the evaluated strains inferred using Mashtree. The tree is based on Mash distances, with the scale bar representing a Mash distance of 0.03 (approximately a 3% difference in average nucleotide identity [ANI]). (**B**) Experimental schematic for data in **C-F**. (**C**) Colony-forming units (CFUs) present on ears at the time of experimental readout. Horizontal bars represent the geometric mean. Dashed line indicates assay limit of detection (30 CFUs). (**D**) Teff (CD62L^−^ CD44^+^) frequencies of CD4^+^ Foxp3^-^ T cells in the ear-draining cervical lymph node (top) and CCR6^+^ Th17 counts in the ears (bottom). (**E**) Teff (CD62L^−^ CD44^+^) frequencies of CD8^+^ T cells in the ear-draining cervical lymph node (top) and Tc17 (CCR6^+^ CD8β^+^) counts in the ears (bottom) after colonization with the indicated strains. Strains that stimulated significant CD8^+^ T cell responses are colored red, all other strains colored blue. (**F**) Per mouse correlation between skin Tc17 count and colonization burden (log10 CFU/ear) across all mice colonized with stimulatory (red) or non-stimulatory (blue) *Staphylococcus* strains (*n* = 98). Tc17 counts were not significantly correlated with colonization burden (Pearson *r* = 0.13, *P* = 0.21, 95% CI -0.073 to 0.32). Dashed line represents a linear fit (shown as a guide). (**G**) PCA of the expression of CCR7, CD40, CD80, and CD86 on 2 subsets of splenic DCs (CD11b^−^ and CD11b^+^) following treatment with heat-killed bacteria, live bacteria, or bacterial conditioned media. Colors denote CD8^+^-stimulatory (red shades) and non-stimulatory (blue shades) *Staphylococcus* strains. Controls are PBS (gray circles), media (gray squares), and cell-stimulatory cocktail PMA/ionomycin (dark gray diamonds). Variance explained by principal components is indicated on the axes. Bar graphs display means ± SD. Flow cytometric data represent at least two experiments with five mice per group. * *P* < 0.05, ** *P* < 0.01, *** *P* < 0.001 (one-way ANOVA with Dunnett’s test relative to naïve conditions).

To explain these distinct CD8^+^ T cell responses, we first hypothesized that CD8^+^-stimulatory strains produce distinct microbe-associated molecular patterns (MAMPs), which activate innate immune receptors, or that non-stimulatory strains produce immunosuppressive molecules, leading to differential activation of dendritic cells (DCs) and, therefore, different T cell priming outcomes. To test this, we treated splenic DCs with live or heat-killed bacteria, or with conditioned media from *Staphylococcus* overnight cultures for different lengths of time and then measured surface expression of canonical activation markers (CCR7, CD40, CD80, CD86). As expected, the DC activation patterns were highly divergent between the pathogen *S. aureus* and commensal *Staphylococcus* strains; however, they did not distinguish CD8^+^-stimulatory from non-stimulatory strains (**Figure 1G**; **Figure S1B**). Thus, skin colonist-specific CD8^+^ responses cannot be explained by differential DC activation, suggesting minimal differences in canonical MAMPs between stimulatory and non-stimulatory strains.

### Bacterial quorum sensing controls the CD8^+^ T cell response to *S. epidermidis* skin colonization

Since DC activation and canonical MAMP expression do not explain differential CD8^+^ T cell activation by *Staphylococcus* colonization, we next took an unbiased and microbe-centric approach to find differences between CD8^+^-stimulatory and non-stimulatory *S. epi* using bulk RNA-sequencing (RNA-seq). Among genes conserved across all six *S. epi* strains, previously described quorum-sensing (QS) effectors comprised 8 out of 10 most upregulated genes in the CD8^+^-stimulatory strains across growth phases (**Figure 2A-B, S2A**), and the upstream regulators of QS were also more highly activated in the stimulatory strains (**Figure 2C**). Thus, we hypothesized that QS activation is required for the CD8^+^ T cell response.

**Figure 2.**
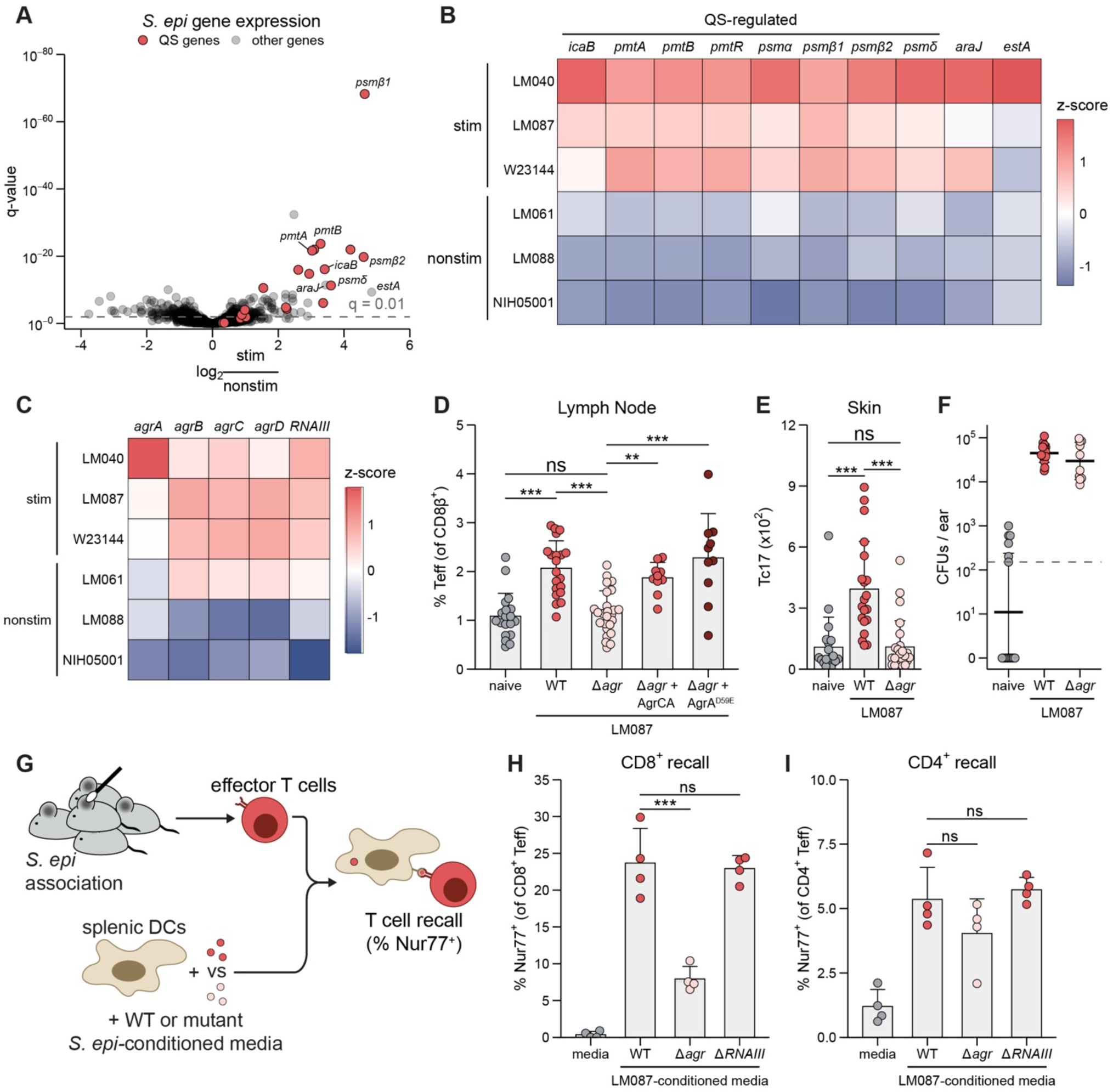
Bacterial quorum sensing controls the CD8^+^ T cell response to *S. epidermidis (S. epi)* skin colonization. (**A**) Differential gene expression between CD8^+^-stimulatory and non-stimulatory *S. epi* strains cultured to exponential phase in Brain Heart Infusion broth (*n* = 3 per strain). Known quorum-sensing (QS) regulators and targets are highlighted in red. Dashed line represents a Benjamini-Hochberg FDR-adjusted p-value (q-value) cutoff of 0.01. Positive log2 fold-changes indicate upregulation in CD8^+^-stimulatory strains. (**B** and **C**) Expression profiles of the 10 most significantly differentially expressed (DE) genes (**B**) and core QS regulators (**C**). Color scales represent batch-corrected, log-transformed mean expression values scaled as per-gene Z-scores. (**D** and **E**) Effector CD8^+^ T cell (Teff; CD62L^−^ CD44^+^) frequencies in the skin-draining cervical lymph nodes (**D**) and Tc17 (CCR6^+^ CD8β^+^) counts in the ear skin (**E**) of naïve controls (gray) or after colonization with the WT (red), Δ*agr* (pink), or Δ*agr* + AgrA^D59E^ (dark red) *S. epi* LM087. (**F**) Ear CFUs at the time of experimental readout. Horizontal bars represent the geometric mean. Dashed line indicates assay limit of detection (150 CFUs). (**G**) Schematic of the T cell recall assay for **H-I**. (**H** and **I**) Reactivation of LM087-elicited CD8^+^ (**H**) and CD4^+^ (**I**) T cells after co-culture with splenic dendritic cells and indicated stimuli, measured by frequency of Nur77^+^ cells. Bar graphs display means ± SD. In vivo experiments show responses pooled from 3 independent experiments. T cell recall data are representative of multiple experiments. * *P* < 0.05, ** *P* < 0.01, *** *P* < 0.001; ns, not significant (one-way ANOVA followed by Tukey’s HSD post-hoc test correcting for all pairwise comparisons; biologically relevant comparisons shown).

QS is a ubiquitous microbe-microbe communication pathway and an important driver of bacterial evolution, predating the divergence between gram-negative and gram-positive bacteria^20^. QS governs cell density-dependent “cooperative” behaviors, such as biofilm maturation and virulence^21^, but its contribution to immune responses during commensal colonization is unexplored. In *Staphylococcus*, QS is controlled by the *agr* (accessory gene regulator) locus, which encodes the sensor kinase AgrC and a master regulator transcription factor AgrA; upon activation, AgrC phosphorylates and activates AgrA, which can directly upregulate hundreds of genes or indirectly regulate genes via RNAIII, a noncoding RNA^22^. To test whether QS is necessary for the CD8^+^ T cell response to *S. epi*, we knocked out both the receptor kinase AgrC and downstream transcription factor AgrA in the stimulatory *S. epi* strain LM087 (Δ*agrCA*, hereafter Δ*agr*). Deletion of QS almost completely abolished the CD8^+^ T cell response in both the lymph node and skin, while leaving colonization largely unchanged (**Figure 2D-F**). Deletion of QS also decreased CD4^+^ T cell responses, suggesting that QS likely provides multiple immune-stimulatory signals, making it essential to all *Staphylococcus* colonization-induced T cell responses (**Figure S2B-C**). Overexpression of either wild-type AgrC plus AgrA or a phosphomimetic mutant of AgrA (AgrA^D59E^)^23^ was sufficient to complement the *agrCA* deletion by restoring the LN CD8^+^ T cell response (**Figure 2D**). Notably, AgrA^D59E^ drove this complementation by elevating QS gene expression on its own in the absence of AgrC (**Figure S2D**). Together with our RNA-seq analysis, these data suggest that higher QS activation is both a natural feature of CD8^+^-stimulatory *Staphylococcus* and necessary for the colonization-induced CD8^+^ T cell response.

Since QS deletion not only reduces CD8^+^ T cell expansion in the skin but also the upstream priming of CD8^+^ Teff in the cLN (**Figure 2D**), we hypothesized that QS controls the expression of either CD8^+^ T cell antigens or auxiliary immunomodulatory molecules required for CD8^+^ T cell priming.

To determine whether QS controls antigen expression, we tested the ability of colonization-induced T cells to be recalled in vitro by wild-type (WT) versus mutant *S. epi*. Based on our initial screen where *S*. *epi* LM087 ear colonization doubled the frequency of CD8^+^ Teff in the cLN (**Figure 1E**), we reasoned that first colonizing mice with LM087 and then isolating CD62L-depleted Teff from the cLN would yield a population that was highly enriched for newly primed LM087-specific T cells. We assessed how well these polyclonal Teff could be recalled by their cognate antigen(s) by measuring the upregulation of Nur77, an early and specific marker of TCR activation^24^ (**Figure 2G**). As expected, splenic DCs co-cultured with LM087-conditioned medium reactivated both CD4^+^ and CD8^+^ Teff while DCs with medium alone did not (**Figure 2H-I**). In comparison to WT LM087, Δ*agr* largely failed to recall CD8^+^ Teff, while Δ*RNAIII* preserved recall, and neither deletion significantly altered CD4^+^ Teff recall (**Figure 2H-I, Figure S2E**). These results suggest that QS is required for the expression of CD8^+^ T cell epitopes and that antigen expression is directly controlled by AgrA and not indirectly by RNAIII.

### The dominant *S. epidermidis* CD8^+^ T cell antigen is an epsilon-class phenol-soluble modulin (PSMε)

To determine the specific QS-regulated genes that provide the CD8^+^ T cell antigens, we took advantage of our finding that QS control of CD8^+^ T cell antigen expression is AgrA-dependent but RNAIII-independent. We found that a class of small, multi-functional proteins known as phenol-soluble modulins (PSMs) are highly AgrA-dependent and less RNAIII-dependent, in agreement with prior literature^25^ (**Figure 3A, Figure S3A**). MS-based proteomic analysis of *S. epi-*conditioned media confirmed that PSM secretion was almost completely abolished in the Δ*agr* mutant (**Figure 3B**). Thus, we reasoned that the PSMs were the most likely candidates for the QS-controlled CD8^+^ T cell antigens.

**Figure 3.**
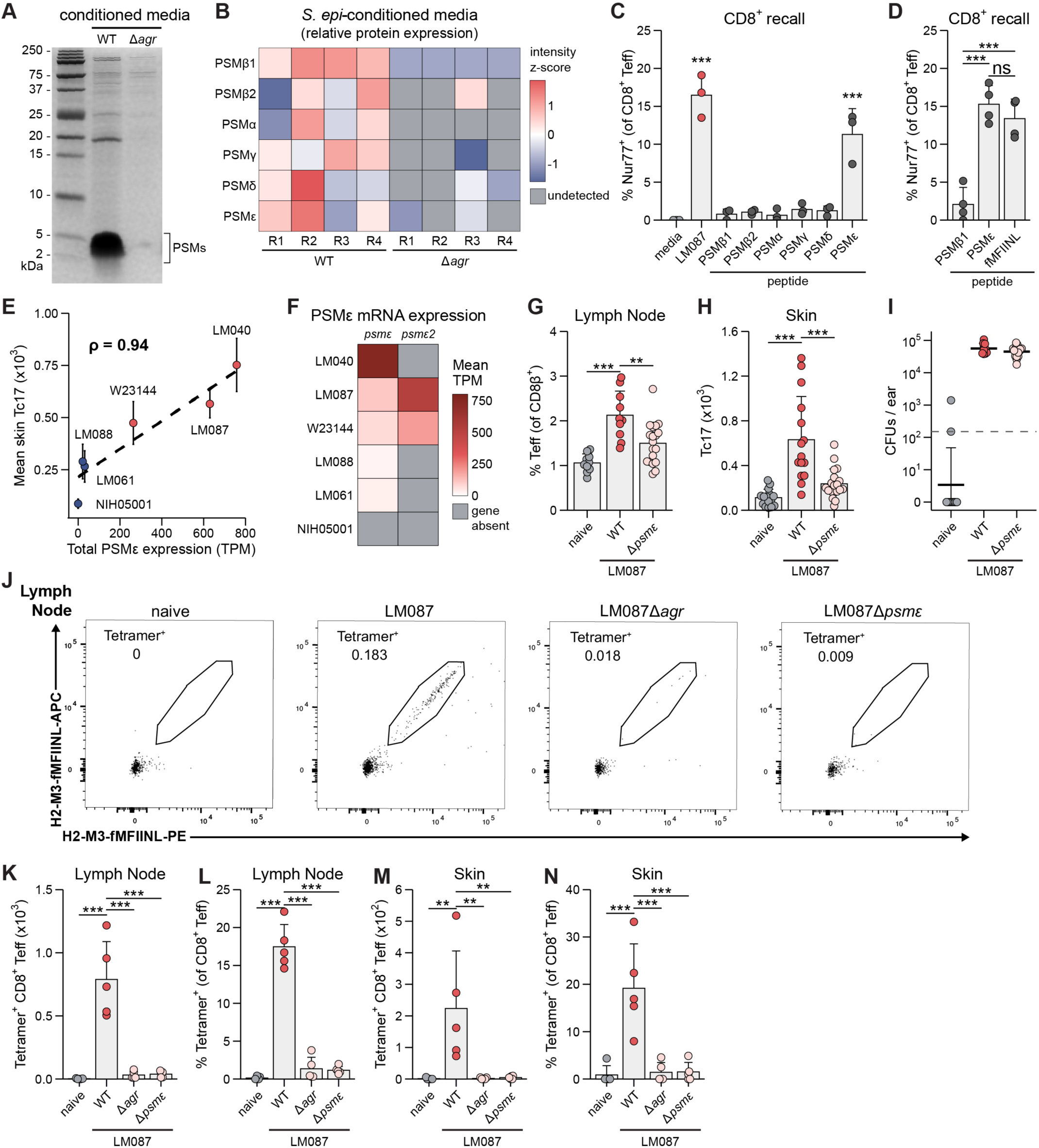
The dominant *S. epidermidis* CD8^+^ T cell antigen is an epsilon-class phenol-soluble modulin (PSMε) (**A**) Coomassie-stained Tris-Tricine gel of *S. epi* LM087 WT and Δ*agr* conditioned media. The prominent band between 2 and 5 kDa is consistent with the expected molecular weight of PSMs (∼2.5 – 4.7 kDa). Molecular weight standards (in kDa) are indicated on the left. (**B**) Relative expression of PSM-derived peptides detected via MS-based proteomic analysis of WT and Δ*agr* conditioned media (*n* = 4 biological replicates per strain). Gray boxes indicate peptides below the limit of detection. Color scale represents normalized, log-transformed values scaled as per-peptide Z-scores. (**C** and **D**) Antigen-specific reactivation (i.e. recall) of LM087 colonization-elicited CD8^+^ T cells after co-culture with splenic dendritic cells and the indicated stimuli (**C**) or synthetic peptides (**D**), as measured by Nur77^+^ expression (PSMγ is also known as δ-toxin; fMFIINL, N-formyl MFIINL). *** *P* < 0.001; ns, not significant (one-way ANOVA with Dunnett’s test relative to the sterile media condition in **C**, and Tukey’s HSD in **D**). (**E**) Correlation between each *S. epi* strain’s total PSMε gene expression (total *psmε* and *psmε2* transcripts per million, TPM) and mean Tc17 count in colonized ears from Figure 1E; error bars represent ± SEM. Spearman *ρ* = 0.94, *P* = 0.016; bootstrap 95% CI ≈ 0.51 – 1.00 (percentile bootstrap, 2,000 resamples). Dashed line represents a linear fit (shown as a guide). (**F**) RNA expression of *psmε* and *psmε2* across each *S. epi* strain. Gray boxes indicate that a given ortholog is not present in a strain’s genome. Color scale represents mean TPM (*n* = 3 biological replicates per strain). (**G** and **H**) Effector CD8^+^ T cell (Teff; CD62L^−^ CD44^+^) frequencies in the skin-draining lymph node (**G**) and Tc17 (CCR6^+^ CD8β^+^) counts in the skin (**H**) of naïve controls (gray) or after colonization with the WT (red) or Δ*psmε* (pink) *S. epi* LM087. (**I**) Ear CFUs at the time of experimental readout. Horizontal bars represent the geometric mean. Dashed line indicates assay limit of detection (150 CFUs). (**J**) Representative fMFIINL:H2-M3 tetramer staining of CD8^+^ Teff in skin-draining lymph nodes from naïve mice or mice associated with WT, Δ*agr*, or Δ*psmε*Δ*psmε2* (Δ*psmε*) LM087. Numbers adjacent to gates indicate fraction of tetramer^+^ cells. (**K**-**N**) Counts and frequencies of fMFIINL:H2-M3 tetramer^+^ CD8β^+^Teff in the skin-draining lymph nodes (**K** and **L**) and skin (**M** and **N**) from naïve mice or mice associated with the indicated LM087 strains. Bar graphs display means ± SD. * *P* < 0.05, ** *P* < 0.01, *** *P* < 0.001 (one-way ANOVA followed by Tukey’s HSD post-hoc test correcting for all pairwise comparisons; biologically relevant comparisons shown).

All *Staphylococcus* express several distinct PSMs, which can vary in number and sequence across strains. *S. epi* typically expresses 6 distinct PSMs, which we obtained as synthetic peptides and tested individually for their ability to recall CD8^+^ Teff elicited by LM087 skin colonization. Only PSMε reactivated CD8^+^ Teff (**Figure 3C**). Additionally, our genome annotation and RNA-seq data revealed that LM087 expresses a homolog of PSMε (PSMε2), which also recalled CD8^+^ Teff (**Figure S3C-D).** Previous work has shown that *S. epi*-specific CD8^+^ T cells are restricted to the nonclassical MHC class Ib molecule H2-M3^8^, which typically presents 6 amino acid epitopes containing an N-terminal formylated methionine^26^, a unique feature of prokaryotic proteins. Thus, we hypothesized that the CD8^+^ T cell epitope was the first 6 amino acids shared by PSMε and PSMε2 (N-formyl MFIINL, i.e. fMFIINL). Indeed, fMFIINL recalled CD8^+^ Teff to the same extent as full-length PSMε (**Figure 3D**), and we confirmed that recall by PSMε or fMFIINL was completely dependent on H2-M3 (**Figure S3F-G**). Supporting its role as the immunodominant antigen, PSMε genome presence and expression levels correlate with in vivo CD8^+^ T cell-stimulatory capacity across *S. epi* strains (**Figure 3E-F**). For example, while stimulatory strains highly express a single copy of PSMε (LM040) or express two copies of PSMε (LM087), non-stimulatory strains minimally express PSMε (LM088, LM061) or lack it entirely in the genome (NIH05001).

Consistent with our T cell recall data, deletion of both PSMε genes from LM087 (Δ*psmε*Δ*psmε2*, hereafter Δ*psmε*) abolished the in vivo CD8^+^ T cell response, despite unchanged colonization load (**Figure 3G-I**). Also, in agreement with our finding that PSMε does not recall *S. epi*-specific CD4^+^ Teff, Δ*psmε* expanded CD4^+^ T cells in the skin-draining lymph node to the same extent as WT (**Figure S3H**). Thus, although QS controls both CD4^+^ and CD8^+^ T cell expansion, the QS-controlled gene PSMε is specifically required for CD8^+^ T cell activation. To further validate PSMε as the immunodominant antigen during *S. epi* colonization, we generated H2-M3 tetramers loaded with the PSMε epitope fMFIINL. In naïve mice, no tetramer-positive cells were detected in the skin and cLN (**Figure 3J-N**). However, 2 weeks after colonization with *S. epi* LM087, 18% of cLN CD8^+^ Teff and 20% of skin CD8^+^ T cells were fMFIINL:H2-M3 tetramer-positive (**Figure 3K**). This percentage is consistent with the percent of LN CD8^+^ Teff recalled in vitro by LM087-conditioned medium (**Figure 3C**). Expansion of these tetramer^+^ CD8^+^ T cells was completely lost when mice were colonized with Δ*agr* or Δ*psmε* (**Figure 3J-N**). The lack of any compensatory CD8^+^ T cell response to alternative antigens suggests that *S. epi* expresses minimal subdominant or cross-reactive epitopes capable of inducing CD8^+^ T cell responses in the setting of commensal colonization.

Together, our in vitro T cell recall, in vivo Δ*psmε* colonization, and tetramer staining data demonstrate that PSMε provides the single immunodominant antigen for CD8^+^ T cells during skin colonization, and its epitope fMFIINL is presented by the nonclassical MHC I molecule H2-M3. Although *S. epi*-specific CD8^+^ T cells are polyclonal, their antigen specificity is exquisitely focused toward a single epitope. Our findings contrast with some infectious models, where H2-M3-restricted T cells have been shown to be cross-reactive. For example, infection with a *Listeria monocytogenes* strain that has the immunodominant epitope deleted leads to unaltered CD8^+^ T cell responses due to high cross-reactivity between multiple H2-M3-restricted antigens^27,28^. Additionally, our RNA-seq and proteomic data demonstrate that the expression of PSMε is tightly controlled by QS activation. Thus, the inability of most of our *Staphylococcus* strains to elicit a CD8^+^ T cell response upon colonization can be explained by their natural loss of PSMε gene content or by their lower QS activation even in the setting of high colonization density.

### PSMε-dependent CD8^+^ T cell activation is conserved across *Staphylococcus*

In our initial screen, *S. cap* also elicited a strong CD8^+^ T cell response upon skin colonization (**Figure 1E**). Given the essential role of QS and PSMε across multiple *S. epi* strains, we hypothesized that *S. cap* also requires QS-activated expression of PSMε for the colonization-induced CD8^+^ T cell response. Consistent with this hypothesis, *S. cap* conditioned media reactivated *S. cap-*induced CD8^+^ Teff, while *S. cap* Δ*agr* did not (**Figure 4A**).

**Figure 4.**
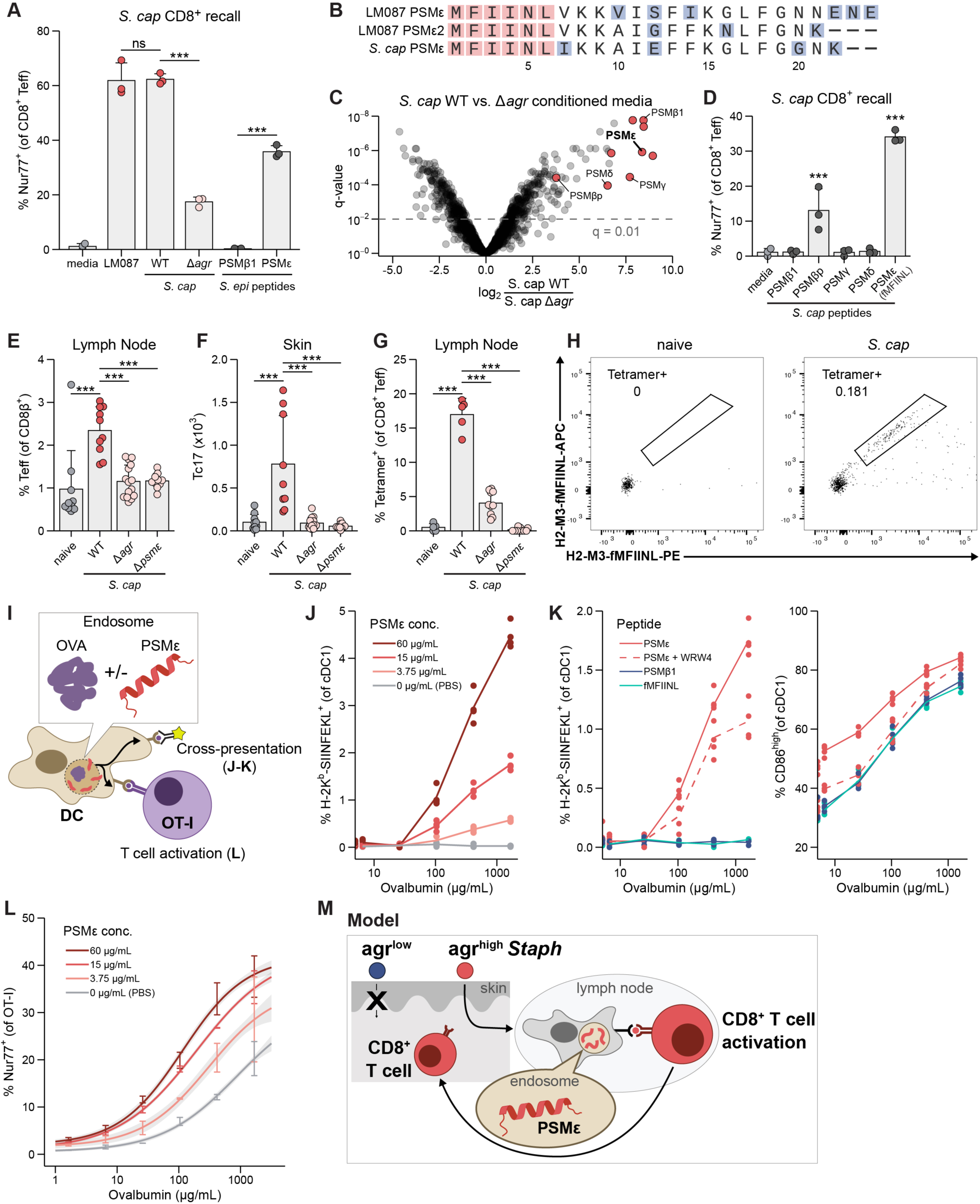
PSMε is the immunodominant CD8^+^ T cell antigen across *Staphylococcus* species and enhances cross-presentation. (**A**) Reactivation of *S. caprae* colonization-elicited CD8^+^ T cells after co-culture with splenic dendritic cells and indicated conditioned media or synthetic *S. epidermidis* (*S. epi*) peptides, measured by Nur77^+^ expression. (**B**) Amino acid sequence alignment of PSMε and PSMε2 from *S. epi* LM087 with the PSMε homolog from *S. caprae* (*S. cap*). The conserved N-formyl MFIINL (fMFIINL) epitope is highlighted in red. Mismatched amino acids are highlighted in blue. (**C**) Differential protein expression in conditioned media from WT and Δ*agr S. cap* cultures (*n* = 4 biological replicates per strain). Red dots indicate predicted PSM homologs. Dashed line represents a Benjamini-Hochberg FDR-adjusted p-value (q-value) cutoff of 0.01. Positive log2 fold-changes indicate upregulation in wild-type. (**D**) Reactivation of *S. cap*-elicited CD8^+^ T cells, performed as in (**A**), using synthetic putative 6 amino-acid N-terminal epitopes derived from the indicated *S. cap* PSMs. As in *S. epi*, the *S. cap* PSMε epitope is fMFIINL. (PSMγ is also known as δ-toxin.) *** *P* < 0.001 (one-way ANOVA with Dunnett’s test relative to sterile media). (**E** and **F**) Effector CD8^+^ T cell (Teff; CD62L^−^ CD44^+^) frequencies in the skin-draining lymph nodes (**E**) and Tc17 (CCR6^+^ CD8β^+^) counts in the skin (**F**) of naïve controls (gray) or after colonization with WT *S. cap* (red) or indicated mutants (pink). (**G**) fMFIINL:H2-M3 tetramer^+^ frequencies of CD8^+^ Teff in the skin-draining lymph nodes of naïve controls (gray) or after colonization with WT *S. cap* (red) or indicated mutants (pink). (**H**) Representative fMFIINL:H2-M3 tetramer staining of CD8^+^ Teff in skin-draining lymph nodes from naïve mice or mice associated with WT *S. cap*. Numbers adjacent to gates indicate fraction of tetramer^+^ cells. (**I**) Schematic of cross-presentation experiments using splenic DCs dosed with exogenous, full-length ovalbumin (OVA) and PSMε. Cross-presentation is observed by directly detecting the MHC-I restricted OVA epitope SIINFEKL presented on H-2K^b^ using an antibody or by measuring Nur77^+^ frequency of OVA-specific OT-I T cells. (**J**) Surface H-2K^b^-SIINFEKL^+^ frequency of cDC1s (CD11c^+^ CD11b^−^ XCR1^+^) after co-incubation with varying concentrations of OVA and increasing doses of PSMε (indicated in legend). (**K**) Surface H-2K^b^-SIINFEKL^+^ frequency (left) and CD86^high^ frequency (right) of cDC1 after co-incubation with varying concentrations of OVA and the corresponding peptide at 15 μg/mL. Dashed line indicates addition of FPR2-antagonist WRW4 (50 μM). (**L**) Nur77 expression in OT-I cells after co-incubation with splenic DCs, varying concentrations of OVA, and increasing doses of PSMε. Error bars represent sample mean ± SEM of data. Lines represent data fitted to a four-parameter log-logistic model. Gray shading around lines represents model prediction ± SE. Bar graphs display means ± SD. * *P* < 0.05, ** *P* < 0.01, *** *P* < 0.001 (one-way ANOVA followed by Tukey’s HSD post-hoc test correcting for all pairwise comparisons; biologically relevant comparisons shown).(**M**) Model: Different species of *Staphylococcus* colonize the skin with different expression levels of quorum sensing (*agr*^low^ or *agr*^high^). *agr*^high^ strains express more PSMε, which both serves as the dominant CD8^+^ T cell antigen and enhances its own antigen presentation in cross-presenting dendritic cells. This drives CD8^+^ T cell priming and subsequent expansion in the skin.

Although *S. epi* and *S. cap* differ by approximately 50% of their gene content (Jaccard index = 0.375), we were surprised to find that *S. cap*-primed CD8^+^ Teff could also be fully recalled by *S. epi*-conditioned media and *S. epi* PSMε, suggesting that *S. cap* shares the same antigen (**Figure 4A**).

However, our initial annotation of the *S. cap* genome identified 8 PSMs but did not yield any clear PSMε homologs. Since small open-reading frames (smORFs) often escape traditional gene finding techniques, we directly searched the *S. cap* genome for potential smORFs with high amino-acid sequence similarity to PSMε using MMseqs2. We identified one such PSMε homolog in the *S. cap* genome, which contains a completely conserved fMFIINL epitope in its N-terminus (**Figure 4B)**. RNA-seq analysis confirmed active transcription of *S. cap* PSMε (**Figure S4A**). Proteomic analysis of the *S. cap* secretome showed that this PSMε homolog is translated and secreted and its expression is lost when QS is deleted (**Figure 4C**). Additionally, in a screen of N-terminal epitopes from each *S. cap* PSM, the PSMε-derived epitope fMFIINL most strongly reactivated *S. cap*-induced CD8^+^ Teff (**Figure 4D)**.

Consistent with our in vitro T cell recall, both QS and QS-controlled PSMε in *S. cap* are required for priming of CD8^+^ Teff in the lymph node and expansion of CD8^+^ T cells in the skin, as both of these responses were lost during *S. cap* Δ*agr* or *S. cap* <u>Δ*psmε*</u> colonization (**Figure 4E-F**). As further confirmation, 20% of cLN CD8^+^ Teff after WT *S. cap* colonization were PSMε-specific, as measured by fMFIINL:H2-M3 tetramer staining (**Figure 4G-H**). Thus, like *S. epi*, *S. cap* expresses a previously unannotated, QS-controlled PSMε homolog that is the immunodominant antigen recognized by *S. cap*-specific CD8^+^ T cells.

### PSMε enhances cross-presentation

We were initially surprised that PSMε was the immunodominant antigen and that this function was conserved across *S. epi* and *S. cap* for two reasons. First, all *Staphylococcus* species express many PSMs, which share high structural similarity (small alpha helix), and all fulfill the H2-M3 preference for N-formylated methionine-containing epitopes, so PSMε did not appear to have more obviously antigenic features compared to other PSMs. Second, PSMε is the least expressed PSM by RNA-seq, over 80-fold lower than the most abundant PSMβ1 in LM087 (**Figure S4B**). Therefore, we hypothesized that, besides harboring the CD8^+^ T cell epitope fMFIINL, PSMε has an additional property that enhances its immunogenicity; specifically, we hypothesized that PSMε increases antigen cross-presentation.

The process of cross-presentation is required to initiate CD8^+^ T cell responses in most contexts; to activate a naïve CD8^+^ T cell, antigen-presenting cells (APCs) must relocate and proteolyze exogenous endosomal contents for presentation on MHC class I molecules, which homeostatically present cytosolic self-peptides^29,30^. Recent studies have found that cross-presentation requires special host factors, such as perforin-2 and WDFY4, which are only expressed within certain APC subsets and promote endosomal leakage of antigens into the cytosol or trafficking of antigens to special vacuoles where they are loaded onto MHC I^31,32^. On the other hand, microbial factors that promote cross-presentation tend to have more nonspecific effects and fall into two categories: PRR ligands that increase APC maturation or virulence factors that help pathogens escape from the endosome into the host cell cytosol as part of an infection^33^. A commensal factor that promotes cross-presentation without infection has not been previously described. However, since PSMs are amphipathic alpha-helices that have been suggested to disrupt cell membranes, we hypothesized that PSMε might facilitate endosomal escape or sorting to cross-presentation-capable vacuoles.

We reasoned that if PSMε could enhance its own cross-presentation through either endosomal escape or endosomal sorting, it should also be able to enhance the cross-presentation of a heterologous antigen. To test this, we incubated splenic DCs with synthetic PSMε and the model antigen ovalbumin (OVA), which must undergo phagocytosis, endosomal escape or sorting, and proteasomal processing before its epitope SIINFEKL can be presented to CD8^+^ T cells (**Figure 4I**). We then directly measured the amount of cross-presentation using a previously validated monoclonal antibody that is specific for H-2K^b^ displaying SIINFEKL^34^. Across a range of OVA concentrations, PSMε, but not PSMβ1, increased H-2K^b^-SIINFEKL presentation by XCR1^+^ classical dendritic cells (cDC1s) in a dose-dependent manner (**Figure 4J-K**). This enhancement of cross-presentation is independent of FPR activation and co-stimulation, as addition of the formyl peptide receptor (FPR) antagonist WRW4 reduced cDC1 expression of the costimulatory molecule CD86 but did not alter PSMε enhancement of H-2K^b^-SIINFEKL presentation (**Figure 4K**). Since we used a mixed population of splenic DCs, we could separately analyze the effect of PSMε on cDC1s versus CD11b^+^ cDC2s. We found that PSMε slightly enhanced cross-presentation in cDC2s but better enhanced H-2K^b^-SIINFEKL presentation in cDC1s (**Figure S4D-E**), which are known to be particularly efficient at cross-presentation^30^. Thus, PSMε enhances cross-presentation via a mechanism that is synergistic with but not completely dependent on pre-existing host factors.

Since PSMε enhanced OVA cross-presentation, we next asked if PSMε could enhance priming of OVA-specific CD8^+^ T cells by co-culturing primary splenic DCs with OVA, naïve SIINFEKL-specific TCR-transgenic T cells (OT-I)^35^, and increasing concentrations of PSMε. PSMε treatment lowered the threshold for OT-I priming in a dose-dependent manner, while a fragment of PSMβ1 did not (**Figure 4L, Figure S4G**). In agreement with our results about the upstream antigen presentation, PSMε enhancement of downstream CD8^+^ T cell priming persisted despite FPR blockade (**Figure S4G**). Consistent with PSMε acting specifically through enhanced cross-presentation, PSMε did not promote OVA-specific CD4^+^ T cell priming, which requires canonical antigen presentation on MHC class II **(Figure S4H).** Thus, PSMε alone is sufficient to promote cross-presentation and subsequent CD8^+^ T cell priming, even to exogenous antigens.

## DISCUSSION

Here, we discover the microbial mechanism underlying CD8^+^ T cell activation by skin-colonizing *Staphylococcus*. We found that PSMε, controlled by QS activation, is required for the CD8^+^ T cell response and acts through a unique mechanism, providing both the antigenic epitope and epitope-agnostic enhancement of cross-presentation. Our findings have both basic science and translational implications.

From a basic science perspective, the importance of *Staphylococcus* QS to the commensal CD8^+^ T cell responses suggests that the mammalian immune system may be “eavesdropping” on a ubiquitous microbe-microbe communication pathway. Notably, PSMε appears uniquely in commensal *Staphylococcus* genomes and has not been found in the pathogen *S. aureus*^36^, raising the possibility that negative selection from host immune pressure has stripped *S. aureus* of PSMε or the alternative possibility that commensals have positively selected for PSMε, which might confer a competitive colonization advantage since the downstream Tc17 response inhibits pathogen growth^1,7^.

Our findings suggest two intriguing future directions. First, because our studies focused on the role of QS activation by a single strain, one future question is how QS interactions among multiple *Staphylococcus* strains shape the CD8^+^ T cell response, as these strains may have cross-strain and cross-species QS synergy or antagonism^37,38^. Since we are colonized by communities of multiple *Staphylococcus* strains, it is possible that antagonistic strains would dampen the T cell response to a strain that is highly stimulatory on its own or that co-colonization by multiple modest QS-activating strains would synergize to generate a robust CD8^+^ T cell response. Second, QS control of T cell antigens may not be limited to *Staphylococcus* since QS is present across the bacterial kingdom and, more specifically, the *agr* system is conserved across the Firmicutes phylum, which includes major gut commensals like *Clostridium* and *Enterococcus*^39^. Therefore, investigating whether QS controls colonization-dependent CD8^+^ T cell responses in the gut and at other barrier sites will be an important future direction.

Additionally, although we focused on QS control of PSMε expression here, it is possible that there are also QS-independent regulators. *Staphylococcus* strains on average harbor 16-17 two-component systems (typically kinase and transcription factor pairs) per genome. These systems convert diverse inputs, such as nutrient availability, ion concentration, and other environmental signals, into gene expression changes. Currently, predicting which two-component systems regulate PSM expression is not possible from the promoter sequence, and many of the inputs that two-component systems sense remain unknown. Our finding that PSMε is a critical factor for host immune responses may now motivate future work to understand its gene regulation, and more broadly to understand how host features and natural strain variation coordinately regulate the immune effects of commensal microbes.

Beyond dissecting the microbial features controlling antigenicity, our findings also advance our conceptual understanding of commensal antigenicity. Though many commensal microbes are now known to induce T cell responses, relatively few have identified T cell antigens. Of those that have been de-orphanized through extensive hybridoma screening, often little of their molecular nature is known outside of their epitope, MHC restriction, and bias for cell surface-associated or secreted proteins^3,4,11,14,40^. We typically think of good T cell epitopes as those that are highly expressed and bind well to MHC. However, PSMε is much more moderately expressed than other PSMs, but has a different distinguishing feature, which is its ability to enhance antigen cross-presentation. Our results suggest that searching for other microbial enhancers of cross-presentation may be a productive way to identify commensals that can activate CD8^+^ T cells upon colonization.

From a translational perspective, our finding that PSMε promotes CD8^+^ T cell activation suggests that it may be useful for vaccination or immunotherapy. Because PSMε promotes antigen cross-presentation without a dependence on PRR engagement and without causing major changes in DC state, it may synergize with existing adjuvants and checkpoint inhibitors. Overall, this study provides a blueprint for dissecting the individual microbial components required for a commensal-specific immune response and then harnessing unique properties of this commensal-immune crosstalk while bypassing live bacterial colonization. Exactly how to best deliver PSMε, whether more potent variants exist, and whether PSMε requires host factors for its function are important questions for future investigation.

## METHODS

### Mice

Wild-type (WT) C57BL/6J, C57BL/6J-CD45a (CD45.1 WT, #002014), and B6.Cg-Tg(TcraTcrb)425Cbn/J (OT-II, #004194) specific pathogen-free (SPF) mice were purchased from Jackson Laboratory. *H2-M3^−/−^* mice were kindly provided by Dr. C. Wang (Northwestern University) via Dr. O. Harrison (Benaroya Institute) and bred in-house with WT C57BL/6J. For in vitro T cell priming, OT-I/*Rag2-/-* mice (#2334) were obtained from Taconic Farms and bred in-house. All mice were bred and maintained under pathogen-free conditions at an American Association for the Accreditation of Laboratory Animal Care (AAALAC)-accredited animal facility at the Broad Institute and housed in accordance with the procedures outlined in the Guide for the Care and Use of Laboratory Animals. All experiments were performed under an animal study proposal (0358-01-23-1) approved by the Broad Institute Animal Care and Use Committee. Sex and age-matched mice between 6 and 12 weeks of age were used for each experiment. When possible, preliminary experiments were performed to determine requirements for sample size, taking into account resources available and ethical, reductionist animal use. Exclusion criteria such as inadequate staining or low cell yield due to technical problems were pre-determined. Animals were assigned randomly to experimental groups.

### Bacterial cultures and topical association

*Staphylococcus* strains were cultured for 18 hours in Brain Heart Infusion broth (BHI) at 37°C shaking^13^. *Corynebacterium* strains were cultured for 18 hours in Brain Heart Infusion broth with 1% Tween-80 (BHIT) at 37°C shaking^41^. *Cutibacterium* strains were cultured for at least 72 hours in BHI at 37°C in an anaerobic chamber, until an OD_600_ of 1.0 was reached. *S. epi* strains LM087, LM088, LM061, LM040 and NIH05001 were obtained from Dr. J. Segre at the NIH^42^. All strains used in this study are listed in **Table S1**. For topical association of bacteria, each mouse was associated by placing 1 mL of the bacterial suspension, approximately 10^9^ colony-forming units/ml (CFU/ml), across the ear skin surface using a sterile polyester swab. Application of bacterial suspension was repeated every 1-2 days for a total of four times. In experiments involving topical application of various bacterial species or strains, cultures were normalized by OD_600_ to achieve similar bacterial density (approximately 10^9^ CFU/mL).

### Tissue processing and flow cytometry

Cells from the ear pinnae were isolated as previously described^1^. Skin-draining cervical lymph nodes were dissociated by mechanical disruption over a 100 μm sterile filter (Miltenyi SmartStrainer) to generate single-cell suspensions. All antibodies are listed in **Table S2** and each staining described below was performed in the presence of purified anti-mouse CD16/32 (clone 93) and 0.2 mg/mL purified rat gamma globulin (Jackson Immunoresearch).

Single cell suspensions were incubated with combinations of fluorophore-conjugated antibodies against surface markers (**Table S2**) in PBS with 5% FBS for 20 min at 4°C and then washed. Fixable Viability Dye eFluor™ 780 (Invitrogen Life Technologies) was used to exclude dead cells.

If tetramer staining was used, PE- and APC-labeled fMFIINL:H2-M3 tetramers (NIH Tetramer Core) were stained at a 1:100 dilution in RPMI with 10% FBS for 30 min at RT before proceeding to surface staining described above.

If intracellular staining was done, after surface staining, cells were then fixed for 1 hour at 4°C using the fixation/permeabilization buffer supplied with the Transcription Factor Staining set (eBioscience 00-5523-00) and then washed. For intracellular staining, cells were then stained with combinations of the following fluorochrome-conjugated antibodies in permeabilization buffer for 1 hour at 4°C: Foxp3 (FJK-16s) and Nur77 (12.14).

Data acquisition was performed on a Cytoflex LX flow cytometer using CytExpert (Beckman Coulter) or LSR Fortessa flow cytometer using FACSDiVa software (BD Biosciences), and data were analyzed using FlowJo software (BD Life Sciences).

Cell counts were calculated using Precision Count Beads (Biolegend) and the manufacturer’s protocol. Cell counts are reported normalized per ear or lymph node.

### Quantification of skin colonization

Skin-colonizing bacteria were isolated at the experiment endpoint by bead-beating one ear from each mouse with sterile 5 mm stainless steel beads (Qiagen #69989) in 1 mL PBS for 5 minutes at maximum speed using a TissueLyser (Qiagen). Serial dilutions were then plated onto BHI or BHIT agar plates and grown at 37°C for 18-48 hours aerobically for *Staphylococcus* and *Corynebacterium* or on BHI agar and grown anaerobically for *Cutibacterium* until single colonies could be clearly discerned by eye to enumerate colony-forming units (CFU). CFUs are reported as normalized per ear.

### Genetic engineering of *Staphylococcus*

Transformation of *S. epi* and *S. cap* was performed as previously described^13^, with the following changes: to generate electrocompetent cells, strains were grown in BHI media with 0.5 M sorbitol (BHIS) (Sigma) and washed in 10% ice-cold glycerol. The resultant electrocompetent cells were transformed with plasmid miniprepped from DC10B *Escherichia coli* by electroporation using a 0.1-cm cuvette (Bio-Rad) and a 1.8 kV pulse with a typical time constant of 2.3-2.5 msec using a Bio-Rad Micropulser. All strains harboring a plasmid were grown in BHI containing chloramphenicol (10 μg/mL).

*S. epi* and *S. cap* allelic replacement was performed as previously described^17^ with temperature-sensitive plasmid pIMAY (gift of Tim Foster, addgene #68939)^43^.

### Design of *Staphylococcus epidermidis* knockout and overexpression constructs

All overexpression constructs were made on the replicative plasmid backbone pLI50-Pcap^13^, derived from pLI50-Ppen-GFP-mut2 or pLI50-Ppen-mcherry (also named pMS182 or pMS183, respectively; gifts from S. Walker, Harvard University). In place of the Ppen promoter, this plasmid features a strong, constitutive Pcap promoter described previously^44^. For AgrA^D59E^ expression, we added the ribosome binding site (RBS) from the *S. aureus* δ-hemolysin gene (*hld*), which promotes strong constitutive translation in *S. epi*^45^. For all other transgenes, each ORF was cloned along with its respective RBS. The AgrA^D59E^ construct was optimized for *S. aureus* using the IDT codon optimization tool. All primers (ordered from Azenta) and synthetic dsDNA fragments (ordered from Twist Bioscience) are included in **Table S1**.

For all overexpression constructs, first, the backbone pLI50-Pcap was linearized with primers oWO_152 and oWO_153, yielding fragment pLI50-Pcap_linearized.

For vector pLI50-Pcap-AgrCA, the *agrCA* CDS was amplified from the *S. epidermidis* LM087 genome using primers oWO_323 and oWO_324, yielding fragment fWO_95. Fragments pLI50-Pcap_linearized and fWO_95 were then assembled via Gibson assembly using the NEBuilder® HiFi DNA Assembly Master Mix (NEB E2621).

For vector pLI50-Pcap-AgrAD59E, fragments pLI50-Pcap_linearized and synthesized fragment fWO_97 (Twist Bioscience) were assembled via Gibson assembly.

To create knockout vectors pIMAY_LM087-agrCA, pIMAY_LM087-RNAIII, pIMAY_LM087-psme, pIMAY_LM087-psme2, pIMAY_Scap-agrCA, and pIMAY_Scap-psme, the following general schema was followed. First, the pIMAY backbone was linearized by PCR with oWO_83 and oWO_84, yielding fragment pIMAY_linearized. Next, the regions upstream and downstream flanking the targeted gene were amplified from the corresponding genome via colony PCR, yielding ∼1 kb upstream and downstream fragments. The fragment pIMAY_linearized, the upstream fragment, and the downstream fragment were then assembled via Gibson assembly.

### High-molecular-weight genomic DNA extraction

For genome sequencing, high-molecular-weight genomic DNA was extracted using the MasterPure Gram Positive DNA Purification Kit (Epicentre, Lucigen). Cell pellets were processed according to the manufacturer’s instructions, with all reagent volumes doubled and vortexing steps omitted to minimize DNA shearing. High-molecular-weight genomic DNA was quantified using a Qubit fluorometer (Thermo Fisher Scientific).

### PacBio single-molecule real-time sequencing and genome assembly

Single-molecule real-time sequencing was carried out on a PacBio Sequel I (Pacific Biosciences) at Fred Hutchinson Cancer Center. Sequencing reads were processed using the Microbial Assembly pipeline within Pacific Biosciences’ SMRT Link v.10.1.0.119588. For *S. caprae*, the resulting assembly comprised two circular contigs of 2,613,786 bp and 45,227 bp.

### Phylogenetics

To infer phylogenetic relationships between strains, assembled nucleotide genomes were analyzed with Mashtree^46^ (version 1.4.6) using default settings.

### Genome annotation and gene clustering

*S. epidermidis* and *S. caprae* genomes were annotated using Bakta^47^ (version 1.11.4) with its “full” database (version 5.0). To identify protein-coding gene clusters across strains, the predicted proteins were clustered using the easy-cluster module of MMseqs2^48^ (version 18-8cc5c) requiring a bidirectional alignment coverage of ≥ 80% and an amino acid identity threshold of 80%. To detect any PSMε homologs missed by Bakta, we queried the *S. epidermidis* LM087 PSMε sequence against the translated genomes using MMseqs2. Hits were filtered to require ≥ 80% bidirectional coverage, a search sensitivity of 7.5, an E-value ≤ 1e-2, and a minimum ORF length of 15 amino acids.

### Bacterial RNA extraction

*Staphylococcus* strains were cultured overnight for 18 hours in BHI at 37°C shaking. Overnight cultures were diluted to an OD_600_ of 0.05 and grown to late exponential phase (OD_600_ of 0.9-1.1), before being collected by centrifugation at 3500g for 8 minutes. Cell pellets were resuspended in 0.5 mL Trizol reagent (ThermoFisher Scientific) and transferred to 2 mL FastPrep tubes (MP Biomedicals) containing 0.1 mm Zirconia/Silica beads (BioSpec Products) and bead-beaten for 90 seconds at 10 m/sec speed using the FastPrep-24 5G (MP Biomedicals). After addition of 200 μL chloroform, each sample tube was mixed thoroughly by inversion, incubated for 3 minutes at room temperature, and spun down 15 minutes at 4°C. The aqueous phase was mixed with an equal volume of 40% ethanol, transferred to a Direct-zol spin plate (Zymo Research), and RNA was extracted according to the Direct-zol protocol (Zymo Research).

### Generation of Bacterial RNA-Seq data

Illumina cDNA libraries were generated using a modified version of the RNAtag-seq protocol^49,50^. Briefly, 250ng of RNA was fragmented, depleted of genomic DNA, dephosphorylated, and ligated to DNA adapters carrying 5’-AN8-3’ barcodes of known sequence with a 5’ phosphate and a 3’ blocking group. Barcoded RNAs were pooled and depleted of rRNA using the riboPOOLS rRNA depletion kit (siTOOLS). Pools of barcoded RNAs were converted to Illumina cDNA libraries in 2 main steps: (i) reverse transcription of the RNA using a primer designed to the constant region of the barcoded adaptor with addition of an adapter to the 3’ end of the cDNA by template switching using SMARTScribe (Clontech) as described^42^; (ii) PCR amplification using indexed primers whose 5’ ends target the constant regions of the 3’ or 5’ adaptors and whose 3’ ends contain the full Illumina P5 or P7 sequences. cDNA libraries were sequenced on an Illumina NovaSeq X platform, generating 150 bp paired-end reads.

### Analysis of RNA-Seq data

Sequencing reads from each sample in a pool were demultiplexed based on their associated barcode sequence using custom scripts (https://github.com/broadinstitute/split_merge_pl). Up to 1 mismatch in the barcode was allowed provided it did not make assignment of the read to a different barcode possible. Barcode sequences were removed from the first read as were terminal G’s from the second read that may have been added by SMARTScribe during template switching. Reads from each sample were aligned to their respective genomes using BWA^51^ and read counts were assigned to genes and other genomic features using custom scripts (https://github.com/broadinstitute/BactRNASeqCount).

Gene expression was compared across related strains using the gene clusters identified above. Differential expression analysis was conducted with DESeq2^52^. Visualization of raw sequencing data and coverage plots in the context of genome sequences and gene annotations was conducted using IGV^53^.

### Reverse Transcription Quantitative PCR (RT-qPCR)

RNA samples (isolated as described above) were analyzed using the Luna Universal One-Step RT-qPCR Kit (NEB #E3005) according to the manufacturer’s instructions, using 2 ng of total RNA per reaction. Amplification was performed on a CFX Connect Real-Time PCR Detection System (Bio-Rad). Data were analyzed using CFX Maestro software (Bio-Rad), and relative mRNA expression (RQ) was calculated using the 2^-ΔΔCq^ method. For *S. epi*, expression levels were normalized relative to *rpoB1*. All reactions performed in technical quadruplicates. All RT-qPCR primers (ordered from Azenta) are detailed in **Table S1**.

### Protein gels and proteomic analysis

Conditioned media from overnight *Staphylococcus* cultures in TSB (protein gel) or minimal media (secretomics) was collected as follows. Bacteria were pelleted at 3500 rcf for 10 minutes, the supernatant was removed, sterile-filtered with a 0.2 μm syringe filter (Cytiva 4612) and concentrated 10x using a 3 kDa MWCO centrifugal filter (Millipore). The resulting concentrate, containing the secretome, was run on a Tris/Tricine gel (Bio-Rad 4563066) and stained with Coomassie Blue for visualization or processed further for mass-spectrometry sample preparation. Protein samples were visualized alongside Precision Plus Protein™ Dual Xtra Prestained Protein Standards (Bio-Rad).

### Mass spectrometry sample preparation

Concentrated bacterial cultures were lysed in SDS lysis buffer (5% SDS, 50 mM TEAB pH 8.5, 2 mM MgCl₂, 2 μg/mL aprotinin, 10 μg/mL leupeptin, 1 mM PMSF) and incubated at 10 °C for 15 min. Samples were supplemented with 3 µL of benzonase nuclease (250 U/µL; Thomas Scientific, E1014-25KU), mixed thoroughly, and incubated for an additional 15 min at 25 °C to degrade nucleic acids.

Insoluble material was pelleted by centrifugation at 20,000 x g for 10 min at 10 °C, and the clarified lysates were transferred to fresh tubes. Protein concentration in the clarified lysates was estimated by BCA assay (Thermo Scientific, # 23227).

Disulfide bonds were reduced with 5 mM dithiothreitol (DTT) for 1 h at 25 °C with shaking at 1,000 rpm, and free cysteines were alkylated with 10 mM iodoacetamide (IAA) for 45 min at 25 °C with shaking at 1,000 rpm in the dark. Lysates were then acidified to a final concentration of 1.2% (*v/v*) phosphoric acid. An aliquot of 50 µg proteins from each sample was precipitated in S-Trap binding buffer (90% methanol, 100 mM TEAB) and loaded onto S-Trap cartridges (Protifi) by centrifugation at 4,000 x g for 1 min, followed by 4x washes with S-Trap binding buffer at 4,000 x g for 1 min. On-cartridge digestion was performed in 50 mM TEAB supplemented with trypsin and Lys-C (each at 1:50 enzyme-to-substrate ratio, *w/w*). The digestion solution was passed through the cartridge by centrifugation at 2,500 x g for 1 min, reapplied to the top of the column, and incubated overnight at 37 °C. Peptides were eluted sequentially with 50 mM TEAB, 0.1% FA, and 2x with 50% ACN/0.1% FA, each elution followed by centrifugation at 4,000 x g for 1 min. Eluates were pooled, snap-frozen, and dried by vacuum centrifugation.

Dried peptides were resuspended in 3% ACN/0.1% FA, and 1% of each sample was loaded onto Evotips and analyzed by LC-MS/MS on a timsTOF HT mass spectrometer coupled to an EvoSep One system.

### Mass spectrometry data analysis

Data from *S. epidermidis* and *S. caprae* (8 samples per species) were analyzed separately in Spectronaut (version 20) using the directDIA+ workflow, with each dataset searched against its corresponding species-specific database. Trypsin/P was specified as the cleavage rule, allowing up to four missed cleavages. Fixed modification was set to Carbamidomethyl (C), and variable modifications were set to Acetyl (Protein N-term), Deamidation (N), FormylMet (Protein N-term), Oxidation (M), with up to five variable modifications allowed per peptide. Identifications were filtered to a precursor Q-value cutoff of 1% and a precursor PEP cutoff of 2%; protein identifications were filtered to experiment-wide and run-wise Q-value cutoffs of 1% and 5%, respectively, and a protein PEP cutoff of 75%. Protein quantities were derived from MS2 peak areas using MaxLFQ, with both cross-run normalization and imputation disabled. All mass spectrometry data is available in **Table S3**.

### Splenic dendritic cell isolation

DCs were isolated from spleens of sex-matched CD45.2 mice that were pretreated with a subcutaneous injection of 1-2×10^6^ Flt3L-producing B16 cells to increase DC yield^54^. The B16-Flt3L cell line was grown in Dulbecco’s modified Eagle’s medium (DMEM, ATCC 30-2002) supplemented with 10% (v/v) fetal bovine serum. Cell growth was maintained at 37°C in a 5% CO2 atmosphere, passaging at a 1:10 dilution every 2-3 days. Mice were sacrificed 10-14 days after injection, and spleens were dissociated using the GentleMACS Spleen Dissociation Kit (Miltenyi 130-095-926). For the DC MAMP screen, untouched DCs were isolated using the Pan Dendritic Cell Isolation Kit (Miltenyi 130-100-875). For DC and T cell co-cultures (i.e. Teff recall and OTI priming), DCs were enriched using CD11c microbeads (Miltenyi 130-125-835) according to manufacturer’s instructions.

### DC MAMP screen

Splenic DCs were isolated as outlined above using the Pan Dendritic Cell Isolation Kit (Miltenyi 130-100-875) according to the manufacturer’s instructions.

1×10^5^ DCs were cultured in DMEM containing 10% FBS with one of the following treatments: PBS, sterile BHI broth, PMA/ionomycin cell stimulation cocktail (eBioscience 00-4970-93), heat-killed bacteria (diluted to ∼10^9^ CFUs/mL in PBS and incubated at 70°C for 30 minutes), live bacteria (in PBS at an MOI of ∼5), or conditioned media from an overnight culture (sterile-filtered with a 0.2 μm syringe filter [Cytiva 4612]). To prevent live bacteria overgrowth in longer experiments, after 3 hours media in all wells was replaced with DMEM containing 10% FBS and 1x Penicillin-Streptomycin (Thermo Fisher 15140163). DCs were incubated for a total incubation time of 3, 8, 18, or 24 hours. After incubation, DCs were stained for CCR7, CD40, CD80, and CD86 surface expression on CD11b^+^ and CD11b^-^ cells. Representative PCA plots from the 3-hour treatment shown in **Figure 1G**. Corresponding flow cytometry data for generating PCA plots in **Figure 1G** are shown in **Figure S1B**.

### In vitro recall of *Staphylococcus*-specific effector T cells

To determine the antigen specificity of polyclonal T cells activated by *S. epi* or *S. cap* colonization, we performed in vitro recall of these T cells by co-culturing them with splenic DCs and conditioned media or peptide antigens as follows.

5×10^5^ CD45.2 DCs (isolated as described above using CD11c microbeads) were co-cultured in DMEM containing 10% FBS (100 μL total volume in a 96-well plate) with one of the following stimulants for 2.5 hours at 37°C in a 5% CO2 incubator: 10 μL of PBS, 1 μL of 1x eBioscience Cell Stimulation Cocktail (Thermo Fisher 00-4975-93), or 10 <u>μL</u> of sterile-filtered conditioned media from overnight cultures. After 2.5 hours, 2.5×10^4^ CD45.1 Teff cells (isolated as described above) were added and co-cultured for another 3.5 hours before surface and intracellular staining (as described above) to measure Nur77 upregulation as a marker of TCR activation.

### Detection of H-2K^b^-SIINFEKL expression

To determine whether PSMε enhances antigen cross-presentation, we cultured splenic DCs (isolated as described above using CD11c microbeads) with the model antigen ovalbumin (OVA) as follows. 2.5×10^5^ DCs were cultured in DMEM containing 10% FBS (100 μL total volume in a 96-well plate) for 6 hours at 37°C in a 5% CO2 incubator with 100 ng/mL – 1.6 mg/mL OVA. Additionally, DCs were treated with one of the following stimulants: 10 μL PBS, 3.75-60 μg/mL PSM peptide (Genscript), and/or FPR antagonist WRW4 (Selleckchem S9818). DCs were then stained with anti-H-2K^b^-SIINFEKL (25-D1.16), anti-XCR1 (ZET), anti-CD86 (PO3), and fixed prior to flow cytometry.

### In vitro priming of OT-I T cells and OT-II T cells

To determine if PSMε enhances antigen priming of CD8^+^ or CD4^+^ T cells, we used the model antigen ovalbumin (OVA) in co-culture with DCs and OVA-specific T cells as follows.

T cells were isolated from spleens and lymph nodes of 8-to-12-week-old SPF CD45.2 OT-I/Rag2-/-mice (CD8^+^ T cells) or OT-II mice (CD4^+^ T cells). DCs were isolated from spleens of sex-matched CD45.1 mice as described above.

1×10^6^ DCs were co-cultured in DMEM containing 10% FBS (100 μL total volume in a 96-well plate) with the following stimulants for 6 hours at 37°C in a 5% CO2 incubator: 10 μL of PBS, 0.1 – 1,677 μg/mL OVA (manufacturer), 3.75 – 60 μg/mL PSM peptide (Genscript), and/or 50 <u>μM</u> of the FPR antagonist WRW4 (Selleckchem S9818). After 2.5 hours, 2.5×10^4^ naïve OT-I or OT-II T cells were added (in separate wells) and co-cultured for the remaining 3.5 hours before surface and intracellular staining (as described above) for Nur77.

### Statistical Analyses

All data processing, visualization and statistical analysis was performed using the software R (version 4.4.1) within the RStudio integrated development environment (version 2025.09.0+387; Posit). Data manipulation and visualization relied primarily on the *tidyverse*^55^ suite of packages (version 2.0.0). In mouse and co-culture experiments, comparison of treatment groups was performed using one-way ANOVA with Tukey’s post-hoc test (comparing all group combinations) or Dunnett’s post-hoc test (comparing treatment groups to a single control). All tests used are indicated in the figure and table legends. *P*-values < 0.05 were considered significant.

For principal component analysis, a model was fitted to the centered, scaled, and log10-transformed geometric mean fluorescence intensity (gMFI) of immune activation markers found in each DC subset using the base R *prcomp* function. Dose-response curves were fitted using a four-parameter log-logistic model from the *drc* package^56^.

## Data and materials availability

The NCBI accession number for genome and RNA sequencing data accession number is pending. All other data needed to evaluate the conclusions in the paper are present in the paper or the supplementary materials.

## SUPPLEMENTARY INFORMATION

Figures S1-4 and Tables S1-S2 are available in the supplementary materials PDF.

Table S3 containing proteomics data is available as a separately downloadable spreadsheet.

## ACKNOWLEDGEMENTS

We are deeply indebted to members of the Chen Lab, R.X., and J.D. for helpful suggestions and comments on the manuscript. We thank Oliver Harrison (Benaroya) and Chyung-Ru Wang for the generous gift of H2-M3^−/−^ mice. RNA-Seq libraries were constructed and sequenced at the Broad Institute of MIT and Harvard by the Microbial ‘Omics Core and Genomics Platform, respectively. The Microbial ‘Omics Core also provided guidance on experimental design and conducted preliminary analysis for all RNA-Seq data. We thank the Broad Institute animal facility staff. This work was supported by an HHMI Hanna H. Gray Fellowship (Y.E.C., C.S.); NIH grant 1R21CA293614-01A1 (Y.E.C., W.O.); Searle Scholars Award (Y.E.C.); John Reed Fund (A.H.); Peter J. Eloranta Fellowship (W.G.); Damon Runyon-Rachleff Innovator Award (Y.E.C., J.J.Y.L., K.L.).

## AUTHOR CONTRIBUTIONS

Y.E.C. and W.O. designed the study, analyzed data, generated the figures, and wrote the manuscript. C.L., N.U., and S.C. performed proteomics and initial analysis. B.B. and J.L. performed RNA-seq and initial analysis. C.D.J. performed genome sequencing of *S. caprae*. W.O., K.L., W.G., A.H., J.J.Y.L., and C.S. performed all other experiments. All co-authors helped to edit the manuscript.

## AUTHOR INFORMATION

Correspondence and requests for materials should be addressed to and will be fulfilled by the Lead Contact, Y. Erin Chen.

