## Supplementary Figures and Tables for "PSMε controls skin commensal CD8^+^ T cell activation"

**This PDF file includes:**

Figures S1-S4

Table S1-S2

Table S3 containing proteomics data is available as a separately downloadable spreadsheet.

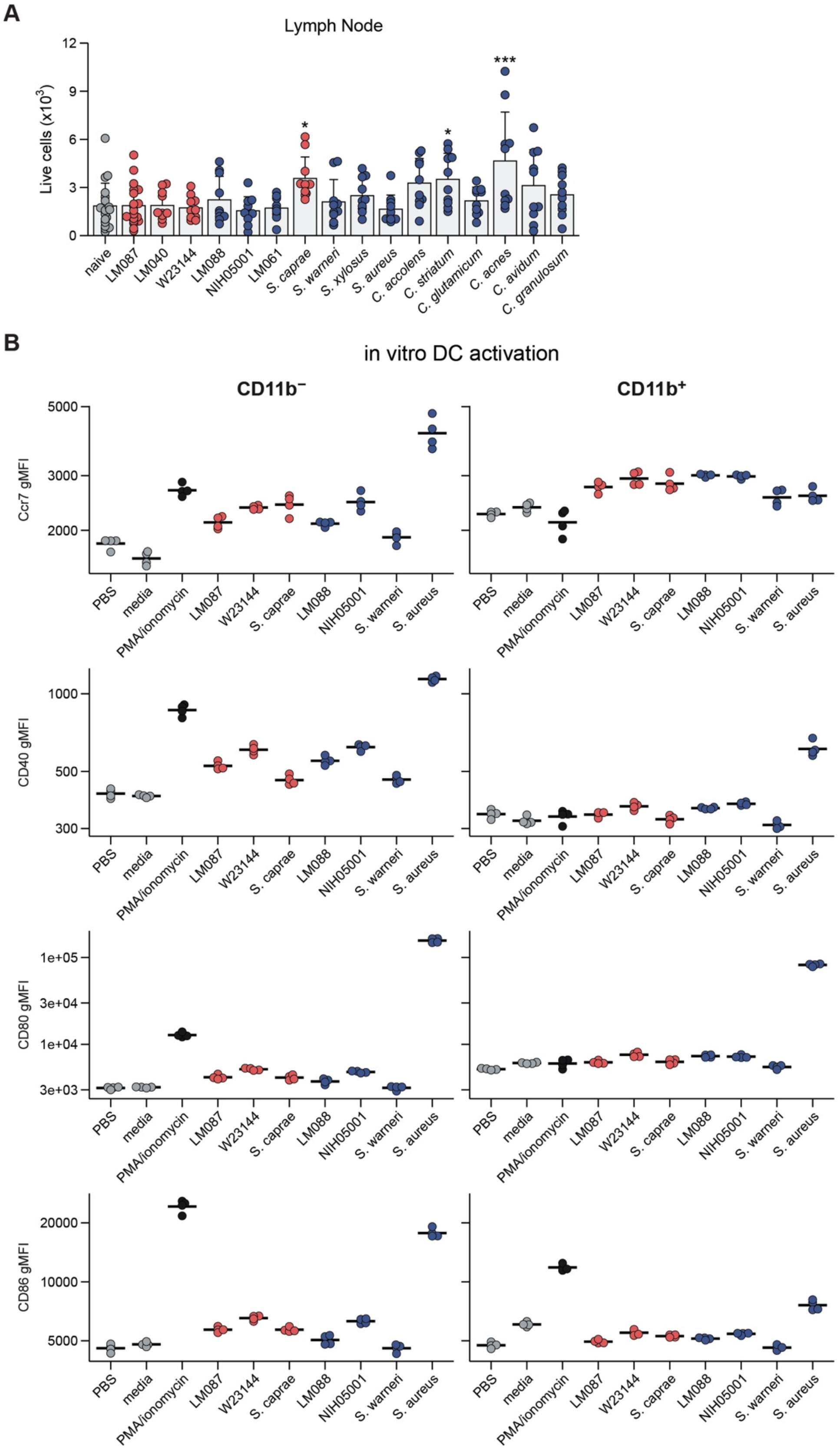

**Figure S1. Supporting data for Figure 1.** (**A**) Total live cell counts in the ear-draining cervical lymph node after colonization with indicated strains. * $P < 0.05$ (one-way ANOVA with Dunnett's test relative to naïve conditions). (**B**) Surface expression of activation markers (CCR7, CD40, CD80, CD86) on CD11b$^-$ (left)

and CD11b$^+$ (right) subpopulations of CD11c-enriched splenocytes after 3-hour treatment with negative controls (gray), PMA/ionomycin positive control (dark gray), live stimulatory strains (red), or live non-stimulatory strains (blue). Live bacteria introduced at a MOI of ~5. Surface expression is reported as geometric mean of fluorescence intensity (gMFI).

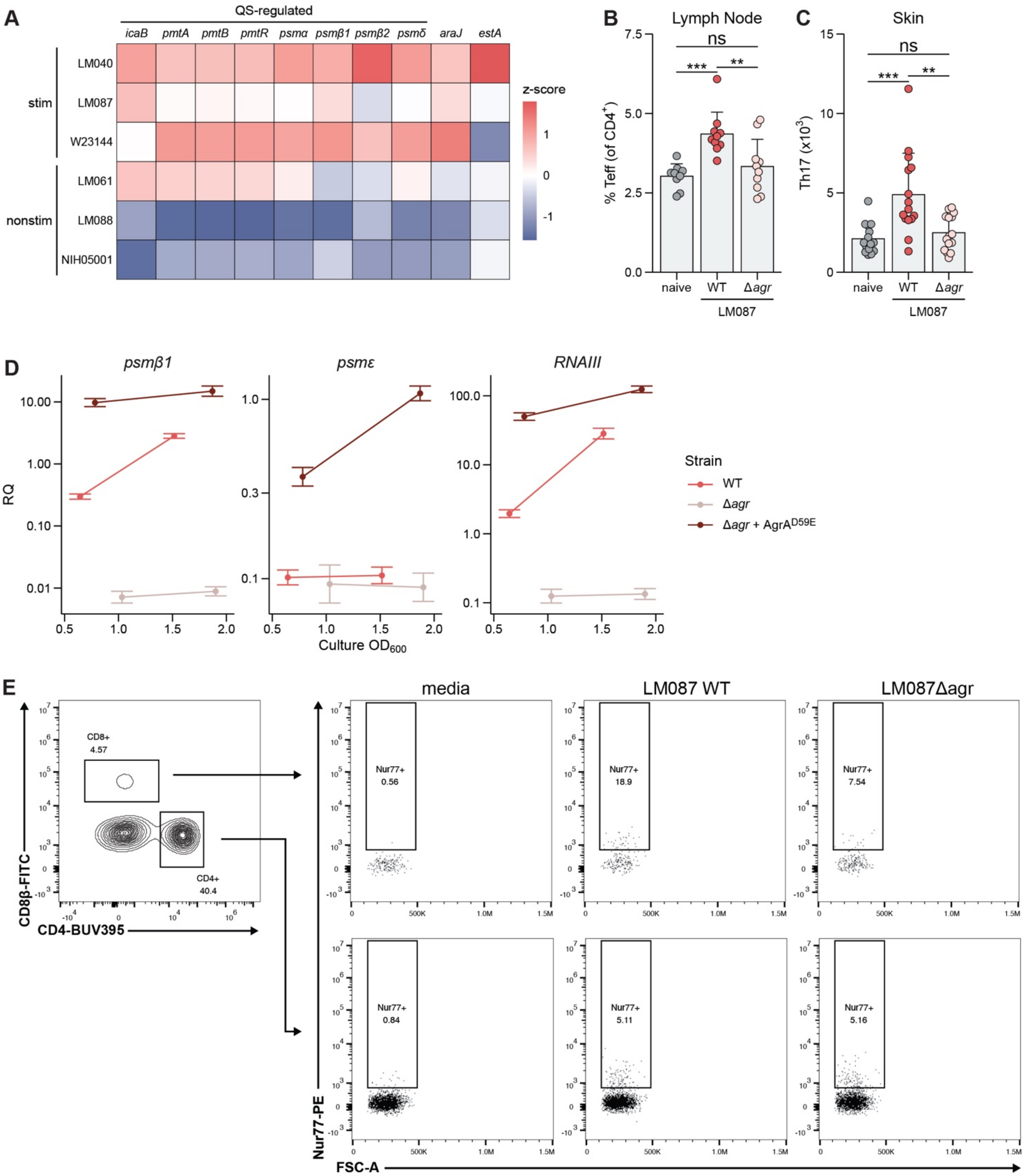

**Figure S2. Supporting data for Figure 2.** (**A**) Stationary phase expression profiles of the 10 most significantly differentially expressed (DE) genes identified in **Figure 2B**. Color scale represents batch-corrected, log-transformed mean expression values scaled as per-gene Z-scores. (**B** and **C**) Effector CD4$^{+}$ T cell (Teff; CD62L$^{-}$ CD44$^{+}$) frequencies in the skin-draining lymph nodes (**B**) and Th17 (CCR6$^{+}$ CD4$^{+}$) counts in the skin (**C**) of naïve controls (gray) or after colonization with the WT (red) or Δ*agr* (pink) *S. epi* LM087. (**D**) Gene expression of *agr* targets in wild-type (red), Δ*agr* (pink), or *Δagr* + AgrA$^{D59E}$ (dark red) strains at culture densities indicated on the x-axis, as measured by RT-qPCR (*n* = 4 technical replicates per sample). RQ, relative quantification (fold change of target relative to housekeeping gene *rpoB1*). Error bars represent mean ± SEM. (**E**) Gating strategy for Nur77 staining in $CD8^+$ (top) and $CD4^+$ (bottom) T cells for **Figure 2H-I**. Numbers inside gates indicate percentage of $Nur77^+$ cells. Bar graphs display means ± SD. * $P < 0.05$, ** $P < 0.01$, *** $P < 0.001$ (one-way ANOVA followed by Tukey's HSD post-hoc test correcting for all pairwise comparisons; biologically relevant comparisons shown).

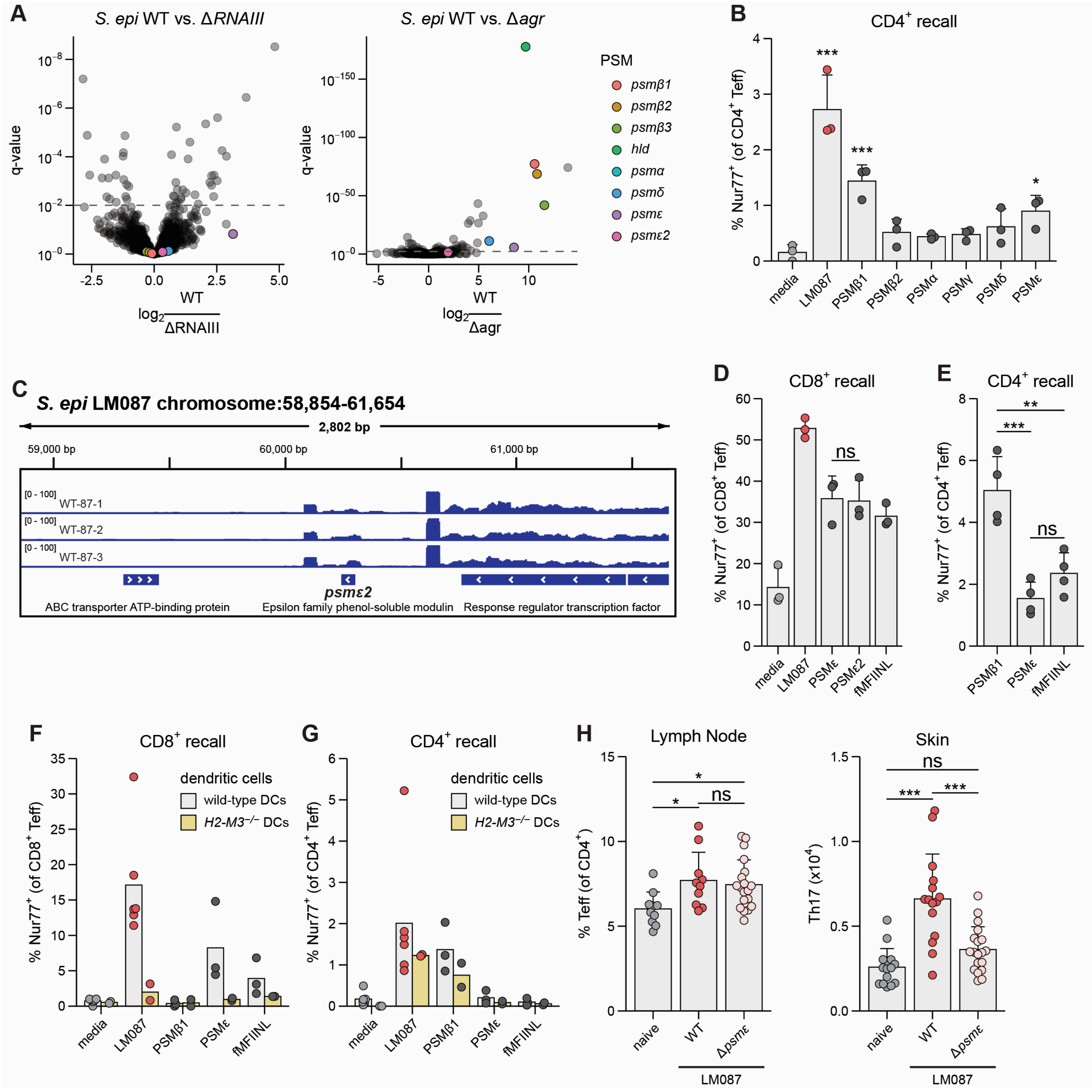

**Figure S3. Supporting data for Figure 3.** (**A**) Differential gene expression between WT and Δ*RNAIII* (left) or WT and Δ*agr S. epi* LM087 (right). PSM genes are highlighted by color according to the legend. Because *hld* is encoded by RNAIII, it is omitted from the Δ*RNAIII* results. Dashed lines represent a Benjamini-Hochberg FDR-adjusted p-value (q-value) cutoff of 0.01. Positive $\log_2$ fold-changes indicate upregulation in the wild-type strains. (**B**) Reactivation of LM087-elicited $CD4^+$ T cells after co-culture with splenic dendritic cells (DCs) and indicated stimuli, as measured by $Nur77^+$ expression. * $P < 0.05$, *** $P < 0.001$ (one-way ANOVA with Dunnett's test relative to sterile media). (**C**) Genomic locus of *psmε2* gene. Track heights represent coverage per million from three biologically independent RNA-seq samples. Gene annotations represent predicted open-reading frames (ORFs). (**D** and **E**) Reactivation of LM087-elicited $CD8^+$ (**D**) or $CD4^+$ (**E**) T cells performed as in (**B**). (**F** and **G**) Reactivation of LM087-elicited $CD8^+$ (**F**) or

CD4$^+$ (**G**) T cells after co-culture with wild-type (gray bars) or *H2-M3*$^{-/-}$ (gold bars) DCs. (**H**) Effector CD4$^+$ T cell (Teff; CD62L$^-$ CD44$^+$) frequencies in the skin-draining lymph nodes (left) and Th17 (CCR6$^+$ CD4$^+$) counts in the skin (right) of naïve controls (gray) or after colonization with the WT (red) or Δ*psmε* (pink) *S. epi* LM087. Bar graphs display means ± SD. * $P < 0.05$, ** $P < 0.01$, *** $P < 0.001$ (one-way ANOVA followed by Tukey's HSD post-hoc test correcting for all pairwise comparisons; biologically relevant comparisons shown).

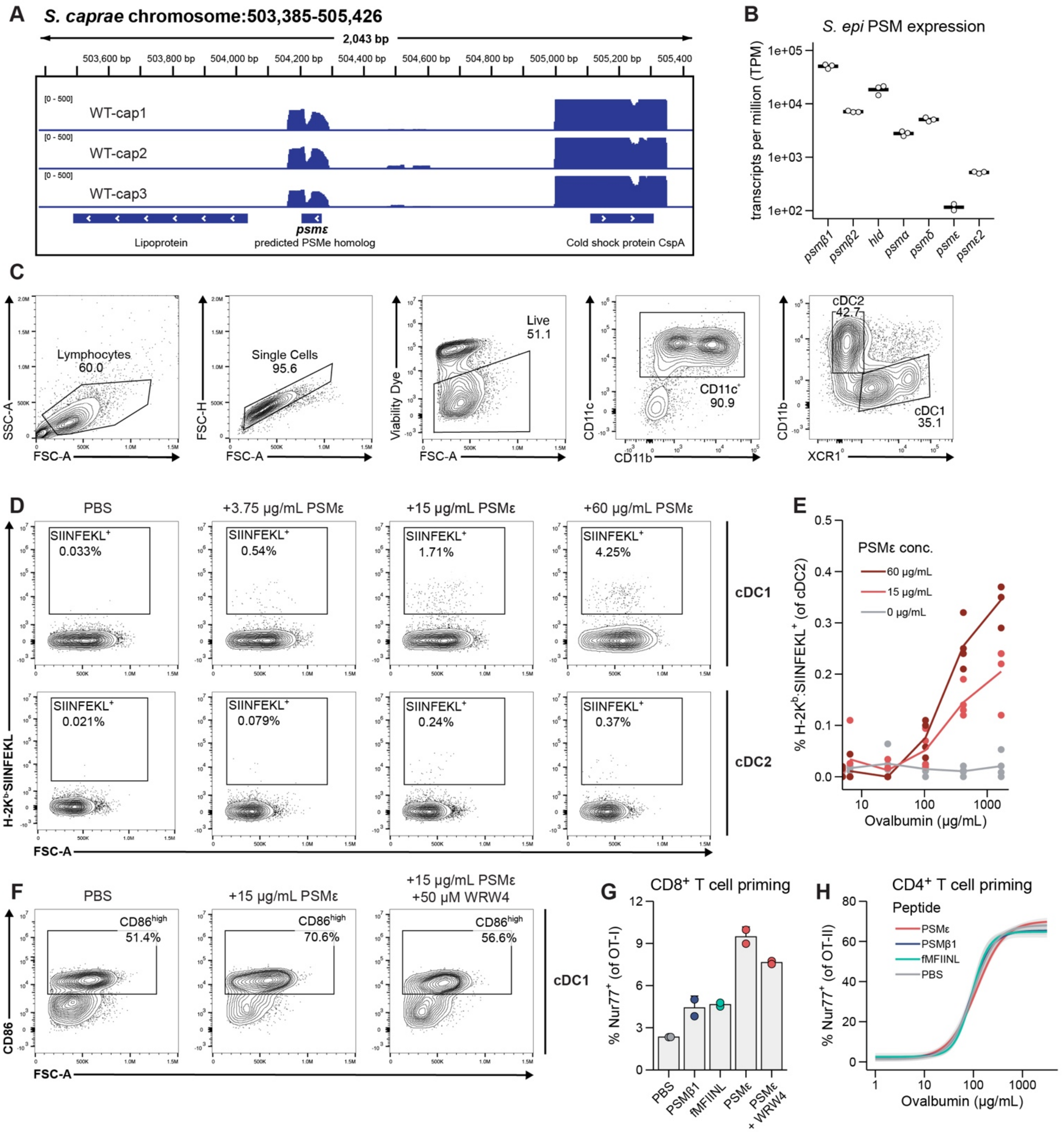

**Figure S4. Supporting data for Figure 4.** (**A**) Genomic locus of *S. cap* PSMε homolog. Track heights represent coverage per million (CPM) from three biologically independent RNA-seq samples. Gene annotations represent predicted open-reading frames (ORFs). (**B**) Gene expression of PSMs in *S. epi* LM087 ($n$ = 3 biological replicates). Bars represent geometric mean of transcripts per million (TPM). (**C**) Gating strategy for dendritic cells. (**D**) Representative H-2K$^b$-SIINFEKL staining of cDC1s (top) and cDC2s (bottom) after incubation with 1.67 mg/mL ovalbumin (OVA) and indicated concentration of PSMε or PBS for **Figure 4J** and **S4E**. (**E**) Surface SIINFEKL$^+$ frequency of cDC2s after co-incubation with varying concentrations of OVA and increasing doses of PSMε (as indicated in legend). (**F**) Representative CD86 staining of cDC1s after treatment with 102 µg/mL OVA and the treatment indicated above each plot. (**G**) Priming of OVA-specific OT-I cells ($CD8^+$ T cells) after co-incubation with splenic DCs, varying concentrations of OVA and corresponding peptide at 15 µg/mL, measured by $Nur77^+$ expression. Dashed line indicates addition of FPR2-antagonist WRW4 (50 µM). Error bars represent sample mean ± SEM of data. Lines represent data fitted to a four-parameter log-logistic model. Gray shading around lines represents model prediction ± SE. (**G**) Priming of OVA-specific OT-II cells ($CD4^+$ T cells) after co-incubation with splenic DCs, peptide, and 26 µg/ml OVA. (**H**) Priming of OVA-specific OT-II cells ($CD4^+$ T cells) after co-incubation with splenic DCs and same conditions as in **Figure 4K**.

| Table S1. Components used for engineering *S. epidermidis* and *S. caprae* | |
|---|---|
| **Primers** | |
| **ID** | **Sequence** |
| oYEC59_RNAIII_fwd | GGCAGCAGATATCATTTCTACAATC |
| oYEC60_RNAIII_rev | GGGATGGCTCAACAACTCAC |
| oYEC99_rpoB1_fwd | gttgtttcagcagcaacagc |
| oYEC100_rpoB1_rev | cgcttgacgttgcatgtttg |
| oYEC103_PSMb1_fwd | AGAAGCAGCCATCACTAACG |
| oYEC104_PSMb1_rev | tcgattcaccatatcaacgctac |
| oWO_345_PSMe_fwd | gttaccgccgaataaacctttg |
| oWO_346_PSMe_rev | aggaggtgagtcaaatgttcatc |
| oWO_83_pIMAY-lin_fwd | gagctccaattcgccctatagtgag |
| oWO_84_pIMAY-lin_rev | caagcttatcgataccgtcgacctc |
| oWO_109_LM087-agrCA_up_fwd | ACGGTATCGATAAGCTTGcgacaagttggtggtgcacc |
| oWO_119_LM087-agrCA_up_rev | CACCTGAGGAGAGTAGgcgaaaattgtttcgcttcagtacg |
| oWO_117_LM087-agrCA_down_fwd | GAAGCGAAACAATTTTCGCCtactctcctcaggtgtcattatacaattttg |
| oWO_118_LM087-agrCA_down_rev | GGCGAATTGGAGCTCgcacagtggcctagctcaag |
| oWO_132_LM087-agrCA_screen_fwd | agacgtctatttgctattggagaggc |
| oWO_133_LM087-agrCA_screen_rev | gacttacgatttccagagatattgcgc |
| oWO_152_Pcap_lin_fwd | tgaaatcactccttccttattttcctttcttg |
| oWO_153_Pcap_lin_rev | ccTATTCTAAATGCATAATAAATACTGATAACATCTTATATTTTG |
| oWO_184_LM087-psme_up_fwd | ACGGTATCGATAAGCTTGcgccgtgcagactttaattttcg |
| oWO_185_LM087-psme_up_rev | GGGCGGAGTAGAGAATAgtgtttgtaatgtagtcgtgtgtgacaaaatg |
| oWO_186_LM087-psme_down_fwd | tattctctactccgcccctcctg |
| oWO_187_LM087-psme_down_rev | GGCGAATTGGAGCTCTAGTggaattatgtatgagggtatcatgagaattata |
| oWO_188_LM087-psme_screen_fwd | cactctgtgagtgtttaacattacttcttgttatcg |
| oWO_189_LM087-psme_screen_rev | gcaagccacctacaaagaactatacatatgg |
| oWO_196_LM087-RNAIII_up_fwd | ACGGTATCGATAAGCTTGctcgtatagtttagtcagttcttctggcac |
| oWO_197_LM087-RNAIII_up_rev | GTGAGCTTGGctttcaatcacatctctgtgatatgagtctattaaaac |
| oWO_198_LM087-RNAIII_down_fwd | TGATTGAAAGccaagctcacggctttaactatgctatttataaag |
| oWO_199_LM087-RNAIII_down_rev | GGCGAATTGGAGCTCTAGTcaaccacctccccattacaatcttttctt |
| oWO_200_LM087-RNAIII_screen_fwd | aagatttgtaggcctgcaaacgg |
| oWO_201_LM087-RNAIII_screen_rev | gatgtagttaccatgaaagcaaagccc |
| oWO_254_LM087-psme2_up_fwd | ACGGTATCGATAAGCTTGgctagcaatgctccagatataggc |
| oWO_255_LM087-psme2_up_rev | AGGTGGCTCACaagagttatcattttcatgtcaaaaaggggtc |
| oWO_256_LM087-psme2_down_fwd | ATAACTCTTgtgagccacctccttgagttatttg |
| oWO_257_LM087-psme2_down_rev | GGCGAATTGGAGCTCTAGTgatcttggtttacctgacatcgatgg |
| oWO_321_LM087-psme2_screen_fwd | ggagtttgatgcagtgtggtcc |
| oWO_322_LM087-psme2_screen_rev | agcgaaagtggggaaagaagc |
| oWO_323-agrCA_fwd | TGGAAAACAAGAAAGGAAAATataaccctagaaagtgtgtaagatatggatg |
| oWO_324-agrCA_rev | AGTATTTATTATGCATTTAGAATAGGcattatattttttaacattacgtac |
| oWG09_Scap-agrCA_up_fwd | CGGTATCGATTAGCTTGccatgtcaacttcactccttccgac |
| oWG06_Scap-agrCA_up_rev | TGGGTTTCATTATGTGTGTAttggtatgtttcaaataattctttaggtactt |
| oWG07_Scap-agrCA_down_fwd | attatttgaaacataccaatacacacataatgaaacccagctttctc |
| oWG08_Scap-agrCA_down_rev | ggcgaattggagctcaggcgcccaaaccaataatgaac |
| oWG_014_agr_screen_fwd | gttgggatggcttaacaatccatttcttag |
| oWG_015_agr_screen_rev | cagagatcctaaagtatttgaaaaggacgg |
| oWG_47_Scap-psme_up_fwd | cggtatcgataagcttgagagagtgaatcattggctaaagctgag |
| oWG_48_Scap-psme_up_rev | aatcctccaattcgtacagaggggcttttcatgcctttataaatattca |
| oWG_49_Scap-psme_down_fwd | atgaaaaagcccctctgtacgaattggaggatttgtcattaaatgcg |
| oWG_50_Scap-psme_down_rev | agggcgaattggagctcatattcgacaaatgatatccccttcgaagc |
| oWG_51_Scap-psme_screen_fwd | gaagcttagcatatgtttatgcggatgg |
| oWG_52_Scap-psme_screen_rev | gtccctgatttagcttccttaatcccc |

| DNA Fragments | |
|---|---|
| **Fragment ID** | **Sequence** |
| fWO_97 | taaggaaggagtgatttcaATGAAAATATTCGTCTGTGAAGACGATCAGAGACAAAGAGAA<br>CACATGGTTTCAATTATAAAAAACTATATTATGATAGAAGAAAAGCCTATGGAATTAGCAT<br>TAGCTACGAATGATCCTTATGAGGTTTTAGAACAATCAAAAGAGTTAAACGATATTGGTTG<br>TTATTTTTTAGAAATTCAATTAGAAGCGGACATGAATGGAATCAAATTGGCGTCAGAGATA<br>AGAAAACATGATCCAGTTGGCAATATTATCTTTGTTACATCACATAGTGAATTAACATATT<br>TGACATTTGTCTATAAGGTCGCAGCTATGGATTTTATTTTTAAAGATGATCCATCAGAGTT<br>AAAAATGCGTATAATTGATTGCTTGGAAACAGCGCACACACGTTTAAAATTATTAAGTAAA<br>GAGTCAAATGTGGATACAATTGAATTAAAAAGAGGCTCAAACTCTGTATATGTGCAATATG<br>ATGATATTATGTTTTTTGAAAGTTCTACAAAGTCTCACCGTTTGATAGCTCATTTAGATAA<br>TCGTCAGATTGAATTTTACGGTAATTTGAAAGAATTAGCACAATTAGATGAGCGTTTTTTT<br>CGTTGCCATAACTCATTCGTGATTAATCGTCATAACATTGAAAGTATTGATTCAAAAGAGA<br>GAATTGTGTATTTTAAAAACGGTGAAAATTGTTTTGCTTCTGTTCGTAATGTAAAGAAAAT<br>TTAAtTATTCTAAATGCATAATAAATACTG |

| Peptides | | |
|---|---|---|
| **ID** | **Sequence** | **Notes** |
| PSMβ1 | fMSKLAEAIANTVKAAQDQDWTKLGT | First 25 amino-acids of *S. epi* LM087 PSMβ1 |
| PSMβ2 | fMEQLFDAIRSVVDAGINQDWSQLAS | First 25 amino-acids of *S. epi* LM087 PSMβ2 |
| PSMα | fMADVIAKIVEIVKGLIDQFTQK | Full-length *S. epi* LM087 PSMα |
| PSMδ | fMSIVSTIIEVVKTIVDIVKKFKK | Full-length *S. epi* LM087 PSMδ |
| PSMε | fMFIINLVKKVISFIKGLFGNNENE | Full-length *S. epi* LM087 PSMε |
| PSMγ | fMAADIISTIGDLVKWIIDTVNKFKK | Full-length *S. epi* LM087 PSMγ, also known as δ-toxin |
| PSMε2 | fMFIINLVKKAIGFFKNLFGNK | Full-length *S. epi* LM087 PSMε2 |
| PSMε_Nterm | fMFIINL | First 6 amino acids of PSMε |
| cap_PSMδ | fMGIVTT | First 6 amino acids of *S. caprae* PSMδ |
| cap_PSMγ | fMASDII | First 6 amino acids of *S. caprae* PSMγ |
| cap_PSMβ1 | fMEKLFD | First 6 amino acids of *S. caprae* PSMβ1 |
| cap_PSMβp | fMSGIVE | First 6 amino acids of *S. caprae* PSMβ homolog |

| Plasmids and strains | | | |
|---|---|---|---|
| **Species** | **Strain** | **Source** | **Other Names** |
| *Staphylococcus epidermidis* | NIHLM087 | Gift of Julie Segre | |
| *Staphylococcus epidermidis* | NIHLM040 | Gift of Julie Segre | |
| *Staphylococcus epidermidis* | W23144 | Gift of Julie Segre | |
| *Staphylococcus epidermidis* | NIHLM088 | Gift of Julie Segre | |
| *Staphylococcus epidermidis* | NIH05001 | Gift of Julie Segre | |
| *Staphylococcus epidermidis* | NIHLM061 | Gift of Julie Segre | |
| *Staphylococcus caprae* | ATCC 55133 | ATCC | |
| *Staphylococcus warneri* | ATCC 27836 | ATCC | |
| *Staphylococcus xylosus* | DBc493 | Gift of Djenet Bousbaine | |
| *Staphylococcus aureus* | NCTC 8325 | Gift of Yasmine Belkaid | |
| *Corynebacterium accolens* | ATCC 49725 | ATCC | |
| *Corynebacterium striatum* | DSM20668 | DSMZ | ATCC 6940 |
| *Corynebacterium glutamicum* | DSM20300 | DSMZ | ATCC 13032 |
| *Cutibacterium acnes* | strain 3 ATCC 29399 | ATCC | |
| *Cutibacterium avidum* | DSM 4901 | DSMZ | KPL1949 |
| *Cutibacterium granulosum* | DSM 20700 | DSMZ | VPI 0507 |

| | | | |
|---|---|---|---|
| *Staphylococcus epidermidis* | LM087Δ*agrCA* | Knockout in NIHLM087 using pIMAY_LM087-agrCA | LM087Δ*agr* |
| *Staphylococcus epidermidis* | LM087Δ*agrCA* / pLI50-Pcap-AgrCA | LM087Δagr transformed with AgrCA overexpression vector | LM087Δ*agr* + AgrCA |
| *Staphylococcus epidermidis* | LM087Δ*agrCA* / pLI50-Pcap-AgrD59E | LM087Δagr transformed with AgrAD59E overexpression vector | LM087Δ*agr* +AgrAD59E |
| *Staphylococcus epidermidis* | LM087Δ*RNAIII* | Knockout in NIHLM087 using pIMAY_LM087-RNAIII | |
| *Staphylococcus epidermidis* | LM087Δ*psme*Δ*psme2* | Serial knockout in NIHLM087 using pIMAY_LM087-psme followed by pIMAY_LM087-psme2 | LM087Δ*psme* |
| *Staphylococcus caprae* | *S. caprae*Δ*agrCA* | Knockout in *S. caprae* using pIMAY_Scap-agrCA | |
| *Staphylococcus caprae* | *S. caprae*Δ*psme* | Knockout in *S. caprae* using pIMAY_Scap-psme | |

| Table S2. Antibodies and dyes used for flow cytometry | | | | | | | |
|---|---|---|---|---|---|---|---|
| **Antigen** | **Species** | **Target species** | **Final Concentration** | **Conjugate** | **Clone** | **Company** | **Catalog #** |
| CD4 | Rat | Mouse | 1 µg / mL | BUV395 | RM-5 | Invitrogen | 363-0042- |
| CD45.1 | Mouse | Mouse | 1 µg / mL | BV421 | A20 | BioLegend | 110732 |
| CD8β | Rat | Mouse | 2.5 µg / mL | FITC | eBioH35-17.2 | Invitrogen | 11-0083-85 |
| Nur77 | Mouse | Mouse | 1 µg / mL | PE | 12.14 | Invitrogen | 12-5965-82 |
| CD45.2 | Mouse | Mouse | 1 µg / mL | APC | 104 | Invitrogen | 17-0454-82 |
| CD90.2 | Rat | Mouse | 1 µg / mL | BV421 | 53-2.1 | BioLegend | 140327 |
| CD8β | Rat | Mouse | 1 µg / mL | BV605 | eBioH35-17.2 | BD Biosciences | 740387 |
| CD62L | Rat | Mouse | 1 µg / mL | FITC | MEL-14 | BioLegend | 104406 |
| TCRβ | Hamster | Mouse | 1 µg / mL | PerCP- | H57-597 | Invitrogen | 45-5961-82 |
| CD44 | Rat | Mouse | 1 µg / mL | PE | IM7 | BioLegend | 103007 |
| CCR6 | Hamster | Mouse | 1 µg / mL | APC | 29-2L17 | BioLegend | 129814 |
| CD11c | Hamster | Mouse | 1 µg / mL | BUV395 | N418 | BioLegend | 744180 |
| XCR1 | Rat | Mouse | 1 µg / mL | BV421 | ZET | BioLegend | 148216 |
| CD86 | Rat | Mouse | 1 µg / mL | BV605 | PO3 | BioLegend | 105125 |
| CD11b | Rat | Mouse | 2.5 µg / mL | FITC | M1/70 | BioLegend | 101205 |
| H-2Kb / | Mouse | Mouse | 1 µg / mL | PE | 25-D1.16 | BioLegend | 141603 |
| H2-Kb | Mouse | Mouse | 1 µg / mL | APC | AF6-88.5 | BioLegend | 116517 |
| CD103 | Rat | Mouse | 1 µg / mL | BUV395 | M290 | BD Biosciences | 568715 |
| CD86 | Rat | Mouse | 1 µg / mL | BV421 | PO3 | BioLegend | 105123 |
| CD40 | Rat | Mouse | 1 µg / mL | BV650 | 3/23 | BioLegend | 124643 |
| CCR7 | Rat | Mouse | 10 µg / mL | PE | 4B12 | BioLegend | 120105 |
| CD80 | Rat | Mouse | 1 µg / mL | PE/Dazzle | 16-10A1 | BioLegend | 104737 |
| CD11c | Hamster | Mouse | 1 µg / mL | APC | N418 | BioLegend | 117309 |
| CD3 | Rat | Mouse | 1 µg / mL | APC-Cy7 | 17A2 | BioLegend | 100221 |
| CD19 | Rat | Mouse | 1 µg / mL | APC-Cy7 | B4 | BioLegend | 115529 |
| CD62L | Rat | Mouse | 1 µg / mL | BV421 | MEL-14 | BioLegend | 104436 |
| TCRβ | Hamster | Mouse | 1 µg / mL | BV711 | H57-597 | BioLegend | 109243 |
| FoxP3 | Rat | Mouse | 2.5 µg / mL | FITC | FJK-16s | eBioscience | 11-5773-82 |
| TCR γ/δ | Hamster | Mouse | 1 µg / mL | PE-Cy7 | GL3 | BioLegend | 118124 |
| CD90.2 | Rat | Mouse | 2.5 µg / mL | AF700 | 53-2.1 | BioLegend | 140323 |
| CCR6 | Hamster | Mouse | 1 µg / mL | BV421 | 29-2L17 | BioLegend | 129818 |
| CD44 | Rat | Mouse | 1 µg / mL | PE-Cy7 | IM7 | BioLegend | 103027 |
| Viability Dye | NA | NA | 1:500 | eFluor-780 | NA | eBioscience | 65-0865-18 |
| CD16/3 | Rat | Mouse | 2.5 µg / mL | NA | 2.4G2 | BD Biosciences | 553141 |
| gamma | Rat | NA | 1 µg / mL | NA | NA | Jackson | 012-000- |
